# SCAF1 driven polyadenylation site usage regulates mRNA isoform expression and neuronal differentiation

**DOI:** 10.64898/2025.12.08.691990

**Authors:** Smaragda Kompocholi, Nikolaos Stamidis, Haiyue Liu, Gertrud M. Hjortø, Eleni Kafkia, Sarah F. Ruidiaz, Thomas C.R. Miller, Jan J. Żylicz, Lea H. Gregersen

## Abstract

Accurate co-transcriptional processing is required for correct gene expression of mRNA transcript isoforms under unperturbed conditions, but particularly during development, to ensure tissue-specific mRNA isoform expression. Here we show that the poorly studied SR-related CTD-associated factors SCAF1 protein regulates polyadenylation site usage towards the end of genes. SCAF1 interacts directly with the phosphorylated C-terminal domain (CTD) of RNA polymerase II (RNAPII), in a complex enriched with elongation and 3’ end processing factors. While *SCAF1* knockout in HEK293 cells is innocuous, it leads to a shift towards expression of shorter mRNA transcripts by co-transcriptional usage of early polyadenylation sites. SCAF1 deficiency induced via auxin-dependent degradation in neuron differentiating mouse embryonic stem cells (mESCs) results in neuronal commitment defects, mediated by altered mRNA isoform usage that impacts expression of key neuronal genes. These findings highlight the importance of mRNA isoform usage and underscores the key role for SCAF proteins in its regulation though polyadenylation site selection.

## Introduction

Transcription is the initial step in the process required for turning genetic information into functional molecules. In higher eukaryotes, most genes encode multiple transcript isoforms, which allow for genomic complexity, making a simple on-off or dosing model of gene expression insufficient to control protein synthesis. mRNA isoform expression is achieved through co-transcriptional mRNA processing including capping, splicing of introns and 3’ end cleavage and polyadenylation ^1,2^. These steps are required not only for proper transcript maturation, but also for gene expression control and transcript isoform regulation, especially during development and differentiation ^3–6^. Since mRNA processing occurs co-transcriptionally, a tight regulation in each phase of transcription is essential for determining the coding and regulatory features of the synthesized mRNA and through inclusion and/or exclusion of exons to secure overall cellular integrity and identity. In terms of mRNA isoform regulation, past work has traditionally focused on alternative splicing; however, it is becoming apparent that mRNA isoform regulation through polyadenylation site (PAS) selection also plays a crucial role ^7–9^. The choice of PAS occurs co-transcriptionally and defines the 3’end of mRNA transcripts: either through usage of early intronic PAS, resulting in mRNAs with different coding potential or through differential usage of PAS within the last exon, leading to mRNA transcripts with distinct 3’UTRs and regulatory potential ^5,10^.

Co-transcriptional processing is orchestrated via the interaction between regulatory factors and RNA polymerase II (RNAPII). To facilitate this, the highly flexible C-terminal (CTD) domain of RNAPII gets extensively modified throughout the transcription cycle ^11–13^, and its phosphorylation status serves as a recruitment platform for mRNA processing factors needed during the different stages of transcription ^14–16^. In a simplified model Ser5 phosphorylated (Ser5P) RNAPII marks early elongation complexes, while Ser2 phosphorylated (Ser2P) RNAPII increases throughout the gene body, with highest levels towards the end of genes ^17^. Accordingly, 5’ capping factors recognize Ser5P RNAPII and 3’ end processing and termination factors, like PCF11, associate primarily with the Ser2 phosphorylated CTD ^13,15,18–20^. In contrast, CTD-binding factors such as mRNA anti-terminators SCAF4 and SCAF8, recognize doubly Ser5P/Ser2P RNAPII ^21^. The importance of proper co-transcriptional mRNA processing is highlighted by the lethal phenotype of the double *SCAF4*/*SCAF8* knockout, due to global shift towards shorter transcript synthesis through increased usage of early intronic PAS ^21^.

Mechanistically, the 3’end processing takes place through a coordinated interplay between RNAPII associated factors, facilitating PAS recognition, and mRNA processing factors. In mammalian cells, 3’end mRNA cleavage is achieved via the action of cleavage and polyadenylation machinery (CPA), consisting of four protein complexes - CPSF (cleavage and polyadenylation specificity factor), CStF (cleavage stimulation factor), CFIm and CFIIm (cleavage factors I and II). Coordinated assembly of these complexes is crucial for regulation of alternative polyadenylation events (APA), with CFI complex enhancing the site-specific assembly of the CPA. PAS usage is differentially regulated during stress and development ^22–25^, allowing for cell adaptation and shaping cell identity through coordinated expression of mRNA isoforms. This is particularly relevant during neuronal differentiation, where many genes undergo regulated lengthening of their 3’UTR ^26–28^. Finally, dysregulation of APA is frequently observed in disease development, such as cancer and neonatal diabetes ^7,8^. Thus, identifying factors involved in APA regulation is essential to understand how PAS selection leading to imbalanced mRNA isoform expression impacts development and disease.

Here we focused on the poorly characterized protein, SCAF1. SCAF1 was originally identified in a yeast-two hybrid screen performed to enrich for interactors of the CTD of RNAPII^29^ and misregulation of *SCAF1* expression has been associated with the development of different types of cancer ^30–33^. It belongs to the family of Serine/Arginine (SR)-CTD-associated factors (SCAFs), along with mRNA anti-terminators SCAF4 and SCAF8 ^21^. SCAF1 is significantly enriched in the interactome of actively elongating RNAPII ^34,35^. Notably, purification of SCAF4- and SCAF8-bound RNAPII captured SCAF1 among the most highly enriched proteins ^34^, showing that SCAF1 interacts with the same RNAPII subcomplexes as SCAF4 and SCAF8.

Through a detailed proteomics and transcriptomics characterization of SCAF1 loss of function models, we show that SCAF1 operates as an mRNA anti-terminator, analogous to SCAF4 and SCAF8. However, while SCAF4 and SCAF8 suppress PAS early in the gene, SCAF1 suppress PAS usage towards the end of genes. Importantly, we show that polyadenylation site suppression by SCAF1 is crucial during neuronal differentiation. Consequently, lack of SCAF1 leads to misregulation of numerous key neuronal genes and defects neuronal commitment.

## Results

### SCAF1 interacts with RNAPII elongation and termination complexes

To assess the cellular localization of SCAF1, we generated stable cell lines expressing GFP-tagged SCAF1. GFP-SCAF1 signal was detected exclusively in the nucleus and localized in nuclear structures in the form of speckles (Fig. 1a). Cellular fractionation using a three-step chromatin extraction by increasing stringency verified the presence of SCAF1 in the nuclease-released fraction (LC), high salt chromatin fraction (HC) as well as the urea (U) fraction, suggesting a tight association of SCAF1 with chromatin (Fig. 1b-c). Of note, through western blots using either an antibody detecting endogenous SCAF1 or epitope-tagged versions of SCAF1 (HA) (Fig. 1c), we consistently observed that the protein migrates slower through PAGE gels than expected (predicted size ∼140 kDa, while SCAF1 migrates at ∼200 kDa). Analysis of the amino acid composition revealed that SCAF1 contain an unusually high percentage of proline residues, which could explain the difference between the predicted and observed migration size (Supplementary Fig. 1a).

**Fig. 1:**
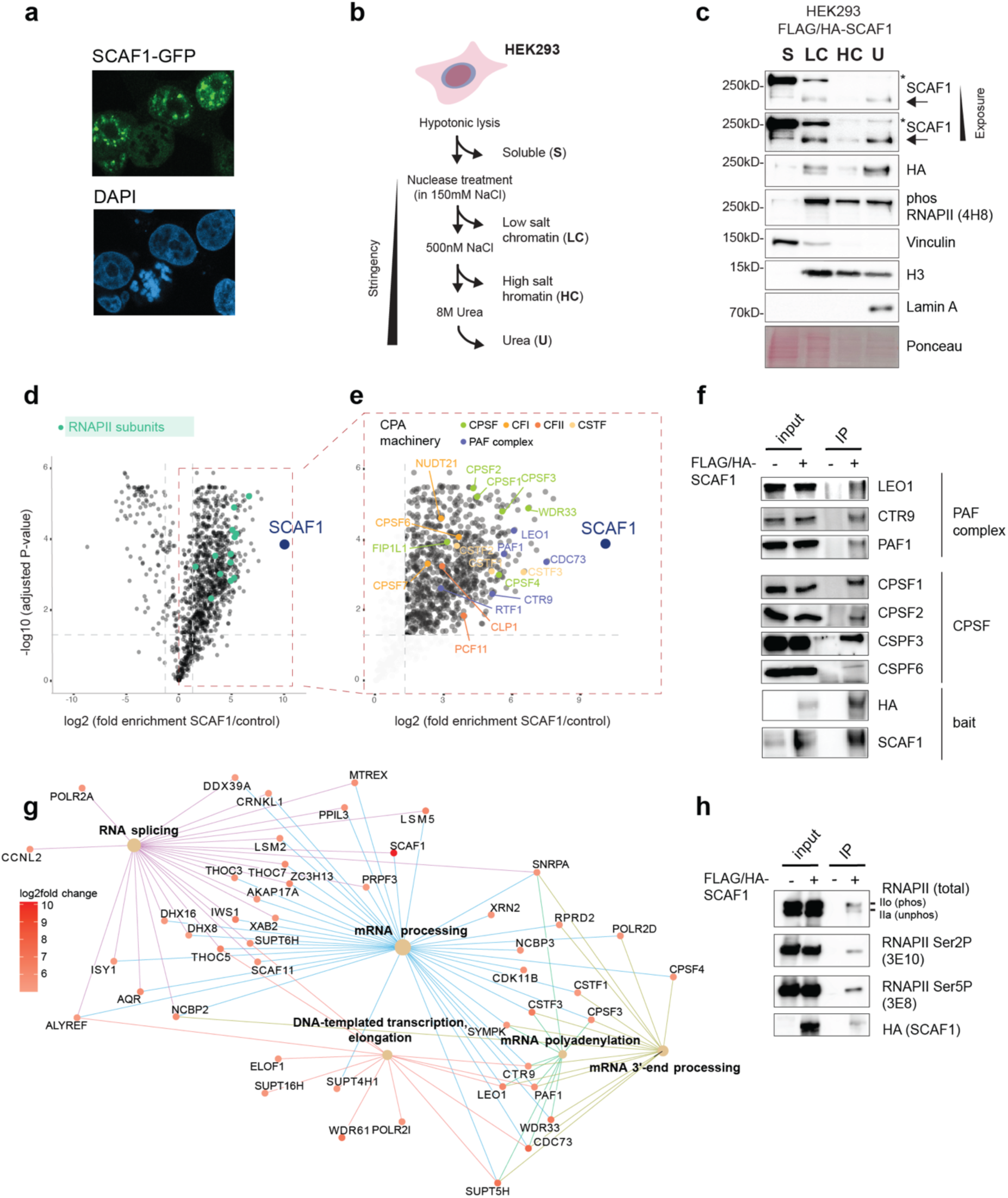
SCAF1 localizes in the nucleus and associates with elongating RNAPII. **a** Cellular localization of stably expressed GFP-SCAF1 in U2OS cells. **b** Outline of cellular fractionation protocol. Fractions from different cellular compartments are isolated, with increasing stringency. Cytoplasmic/soluble fractions (**S**), and chromatin-associated fractions of different strength, DNase/RNase released proteins, low chromatin bound (**LC**), high salt released bound to chromatin fraction (**HC**) and proteins only soluble by 8M urea (**U**). **c** Western blot for protein abundance in the different cellular fractions, isolated as shown in (B). Urea (**U**) is added in the final fraction for isolating the remaining proteins. Proteins were extracted from HEK293 cells stably expressing FLAGHA-SCAF1. Vinculin is used as an internal control for soluble fractions (S), whereas Histone 3 and Lamin A are used for internal controls of chromatin fractions. Two different exposures of SCAF1 blots are shown. Arrows indicate SCAF1 band and asterisk indicates the presence of non-specific bands (confirmed by analysis of *SCAF1* KOs). **d, e** Proteomics of FLAG immunoprecipitates of FLAGHA-SCAF1 expressing HEK293 cells. Y axis indicates the Log2 fold change of the SCAF1 associating proteins, compared to control HEK293 cells not expressing any epitope-tagged proteins. In (**d**) the subunits of RNAPII are highlighted and in (**e**), PAF1 complex components and the components of the four core 3’ end processing complexes are highlighted. **f** Western blot validation of the most highly enriched proteins in FLAGHA-SCAF1 interactome. SCAF1 and HA probing was used to detect the FLAGHA-SCAF1 bait protein. **g** Cnetplot visualization showing top 5 enriched functional categories in color coded edges identified from enriched (log2FC >6) proteins associated with SCAF1, through proteomics data analysis. Proteins are represented by dots and are colored based on their log2 fold change. **h** Western blot of phosphorylation-specific RNAPII-CTD forms in FLAGHA-SCAF1 immunoprecipitants. Antibody recognizing the N-terminal Rpb1 (D8L4Y) was used to detect total RNAPII.

To analyze the SCAF1 interactome, we performed proteomics analysis of SCAF1 immunoprecipitates from chromatin-enriched fractions of HEK293 cells stably expressing epitope tagged SCAF1 (FLAGHA-SCAF1). We confirmed that SCAF1 associates strongly with all RNAPII subunits (Fig. d) and numerous RNAPII-bound proteins. Amongst the highest scoring SCAF1 interactors we found all components of the PAF1 complex (PAF1, LEO1, CDC73, CTR9, RTF1), as well as CPA factors (all components of CPSF, CStF, CFIm and CFIIm) (Fig. 1e and Supplementary Table S1). The interaction of SCAF1 with both PAF complex components, as well as individual CPA components were verified by immunoprecipitation (IP), followed by western blot (Fig. 1f). Moreover, protein components of various complexes involved in co-transcriptional processing and mRNA regulation were also highly enriched in the SCAF1 proteome, most likely mediated through the interaction with SCAF1-bound RNAPII, rather than being direct SCAF1 interactors (Fig. 1g, Supplementary Fig. 1b-c). Using phosphorylation specific RNAPII CTD antibodies, we found that SCAF1 interacts with the hyperphosphorylated, transcriptionally engaged form of RNAPII, including both Ser5P, Ser7P and Ser2P RNAPII (Fig. 1h, Supplementary Fig. 1c). Taken together, these data show that SCAF1 binds RNAPII engaged in active transcription and associates with key RNAPII-bound factors involved in elongation and 3’end mRNA processing.

### Loss of SCAF1 results in increased usage of proximal polyadenylation sites and expression of shorter mRNA isoforms

To address the role of SCAF1 in transcription, we performed mRNA sequencing (mRNA-seq) of two SCAF1 doxycycline-inducible rescue cell lines in *SCAF1* knockout (KO) backgrounds (Fig. 2a-b). Notably, lack of SCAF1 did not markedly affect cell growth or viability in the time scales used for the transcriptomics experiments (Supplementary Fig. 2a). To investigate changes in co-transcriptional processing upon loss of SCAF1, we analyzed the mRNA-seq data using rMATs ^36^, focusing on alternative splicing events. Of the different splicing event types, *SCAF1* deletion significantly increases exon skipping (Fig. 2c). To further identify the positions of the differentially used exons affected by loss of SCAF1, we computed the relative position of individual exons that were differentially expressed in *SCAF1* KO versus WT cells, using DEXseq analysis ^37^. We found 853 genes containing exons differentially expressed in a SCAF1 dependent manner distributing throughout the gene body of the mRNA transcripts. Remarkably, there was a clear trend towards the last exon being most frequently affected in *SCAF1* KO cells (Fig. 2d). Considering the enrichment of CPA factors in the SCAF1 interactome, we next questioned whether SCAF1 might affect splicing through the choice of polyadenylation sites, similar to what has previously been described for SCAF4 and SCAF8 ^21^. To directly access whether SCAF1 affects APA site choice, we used APAlyzer ^38^ to quantify differences in the usage of annotated APA sites. Indeed, SCAF1 loss resulted in differential usage of intronic APA sites in 265 genes (Fig. 2e). Out of the 265 genes, 168 displayed upregulated usage of intronic APA. Comparing the affected genes identified by DEXseq and APAlyzer, we found a subset of 128 genes shared between the two (Supplementary Fig. 2b), which likely represent a conservative estimate of SCAF1 targets, as APAlyzer only considers a predefined set of annotated APA events. Further inspection of individual genes with increased usage of intronic polyA sites confirmed upregulation of mRNA transcript isoforms corresponding to usage of a more proximal annotated polyadenylation site (Fig. 2f-g, Supplementary Fig. 2c-e). This was further validated by qPCR using primer pairs targeting the 3’UTR of the short and long mRNA transcript variant, respectively (Fig. 2f, Supplementary Fig. 2c). Interestingly, specifically for USP15, the increased expression of the shorter isoforms also led to the markedly increased synthesis of a shorter, truncated version of the annotated protein (Fig. 2h). In fact, the shorter protein accumulated in the cytoplasm, accompanied by a subsequent reduction of the nuclear USP15 (Fig. 2h).

**Fig. 2:**
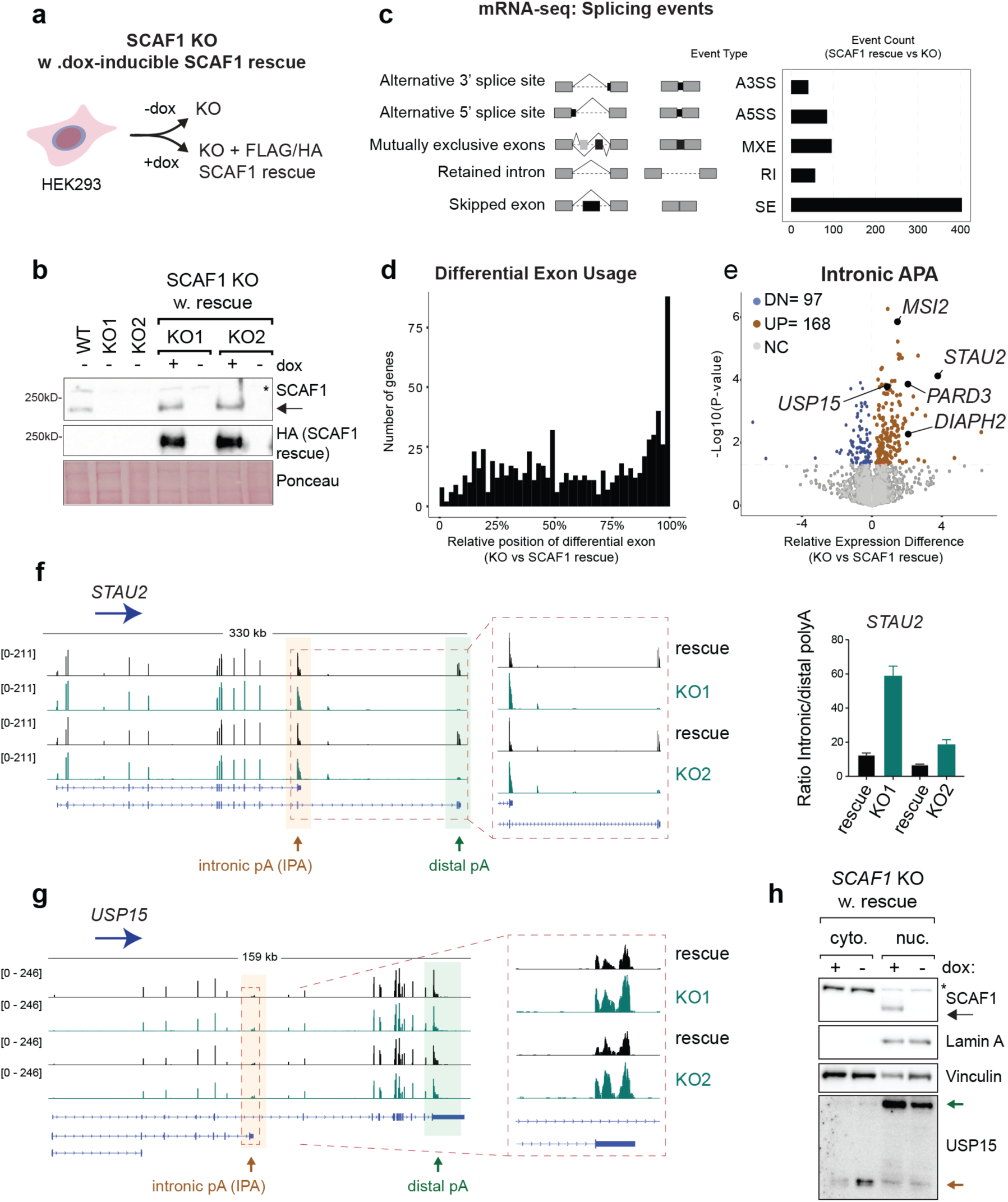
Loss of SCAF1 drives alternative polyadenylation of target transcripts. **a** Scheme for the generation of the different cell lines used for transcriptomics. Individual HEK293 CRISPR Cas9 *SCAF1* KO clones were used as background genotype and were re-introduced epitope-tagged SCAF1, referred to as KO (-dox) and WT/rescue cell lines (+dox) respectively. **b** Western blot validation of the different cell lines used (*SCAF1* KO and doxycycline inducible rescue) in two clones tested. Asterisk indicates non-specific bands and arrow SCAF1. **c** rMATS analysis with graphical representation of alternative splicing events and barplot displaying the number of differential splicing events found as significantly different between samples with and without SCAF1 expression (two individual clones, each grown and harvested in triplicates), based on the mRNA-seq data. **d** Histogram displaying the relative position of differential exon usage, upon loss of SCAF1 (DEX-seq analysis). **e** Volcano plot, generated after APAlyzer analysis, showing differentially used intronic alternative polyadenylation (APA) sites in *SCAF1* KO, compared to rescue conditions. Data were merged from two clones, sequenced in triplicates per SCAF1 genotype background. Colored dots represent transcripts with significantly (p-value <0.05) increased (dark red) or decreased (blue) usage of the annotated intronic APA sites. **f, g** IGV tracks displaying mRNA-seq results from two *SCAF1* KO clones tested (KO1 and KO2) and their corresponding rescue samples. Orange and green arrows indicate the annotated sites of intronic and distal polyadenylation (polyA) sites respectively. **f** Reduced read signal of the long mRNA isoform, validated by qPCR. Primer-pairs used for detecting the two isoforms were designed to be specific for the short isoform 3’ UTR or the long isoform 3’ UTR. The ratio of intronic to distal polyA was calculated as the ratio between products from the primers targeting the short isoform 3’ UTR to long isoform 3’ UTR, normalized to *GAPDH* and a gene specific reference region (intron spanning primer pair common for both isoforms). Error bars represent ±SD. **h** Western blot using USP15 antibody against the N-terminus of the protein, able to detect full length USP15 (green arrow) and short form (orange arrow) in fractionated samples of SCAF1 rescue and KO. SCAF1 signal is indicated with the black arrow and asterisk marks non-specific bands.

Collectively, these findings reveal that lack of SCAF1 results in usage of more proximal polyA sites, leading to shorter mRNA isoforms being favored over the longer mRNA isoform for a specific subset of genes in HEK293 cells.

### Isoform switch events controlled by SCAF1 occur co-transcriptionally

To determine whether the choice of polyadenylation site happens co-transcriptionally or is a post-transcriptional effect of SCAF1 on the stability of the different isoforms, we performed nascent RNA sequencing using TT_chem_-seq to capture RNAPII activity across genes ^39^ (Supplementary Fig. 3a). In parallel, we captured newly synthesized, full-length mRNA transcripts by labelling RNA with 4-thiouridine (4sU) for 60 min, followed by mRNA-seq (4sUmRNA-seq) (Supplementary Fig. 3b). This allows us to look at full-length mRNA transcripts produced exclusively during the 4sU labelling period and thus limiting the effect of mRNA stability. First, we confirmed that the SCAF1-dependent changes in mRNA isoform expression identified from steady-state mRNA-seq analysis was recapitulated in the 4sUmRNA-seq, indicating that the changes indeed happen co-transcriptionally and are not due to changes in mRNA stability (Fig. 3a-b). 3’end mRNA cleavage and RNAPII termination is a two-step process, first triggered by mRNA cleavage downstream of the polyadenylation signal which generates the 3’end of the mature transcript; this subsequently leads to transcription termination, where RNAPII disassociates from the DNA template, typically after transcribing an additional 4-10 kb region downstream of the cleavage site ^2,17,40^. To assess whether upregulated early mRNA cleavage events also result in early RNAPII termination upon loss of SCAF1, we analyzed the TT_chem_-seq signal between the proximal and distal polyadenylation sites. Single gene examples such as *STAU2* and *PARD3*, showed that the TT_chem_-seq signal indeed decreased in both *SCAF1* KO clones between the proximal and distal polyadenylation sites (Fig. 3a-b), indicating that RNAPII indeed prematurely terminates downstream of the proximal polyadenylation sites in these cells. The same trend was recapitulated in the subset of genes displaying upregulated intronic APA site usage (168 genes; Fig. 3c). Close examination of the area near the annotated transcription end site (TES), showed that that the TT_chem_-seq signal indeed significantly drops in both *SCAF1* KO clones (Fig. 3d). Reassuringly, other than close to the TES, we do not observe significant differences in the signal density anywhere else in the gene body of those targets (Supplementary Fig. 3f). Moreover, no overall defect in transcription dynamics or levels was observed, assessed with both genome wide nascent transcriptomics (TT_chem_-seq signal) and imaged based quantification of the newly synthesized transcripts (EU labelling), upon deletion or depletion of SCAF1 respectively (Supplementary Fig. 3c-e). This further supports the specificity of SCAF1 controlling the polyadenylation site of a defined subset of target genes co-transcriptionally, resulting in subsequent early RNAPII termination of a specific gene set rather than genome-wide effects on RNAPII dynamics.

**Fig. 3:**
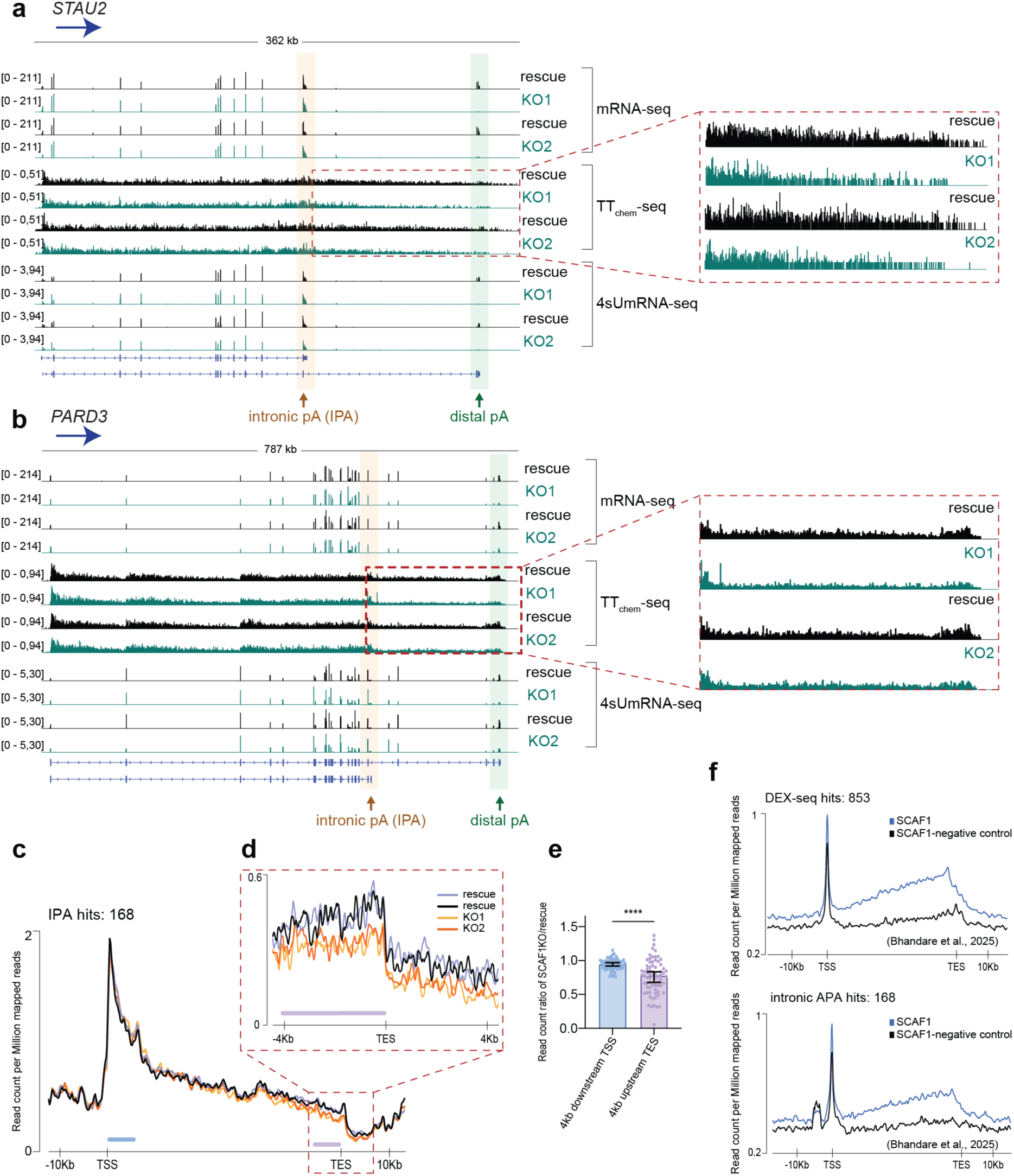
SCAF1 acts co-transcriptionally and suppresses premature termination of target genes. **a, b** IGV tracks, comparing the data from the different RNA sequencing methods. Orange and green arrows indicate intronic and distal polyA sites respectively. **c** Metagene profile of TT_chem_-seq, against the 168 target genes of SCAF1 that display increased usage of intronic polyA sites. The two SCAF1 clones are presented (with merged duplicates) and gene size is scale normalized with 10 kb region upstream and downstream of the genes plotted. **d** Zoom in around the transcript end site (TES) of the metagene plot shown in (**c**). Plot centered to TES, with 4kb upstream and downstream presented. **e** Read coverage quantification of the TT_chem_-seq data presented in (**c**). Read count (RPKM) ratio of the signal from the *SCAF1* KO, compared to rescue (y axis), measured in a 4 kb window upstream from TES and in a 4 kb window downstream of TSS for the 168 upregulated IPA genes (168). Calculations were performed using build in Easeq tools. Statistical significance measured by running Wilcoxon test (p-value <0.0001, ****). **f** Metagene analysis of SCAF1 CUT&RUN data from ^41^ for HA-tagged SCAF1 in U2OS cells. Read densities averaged over SCAF1 target genes either with differential exon usage analysis (DEX-seq) (upper part of the panel, 853 genes) or displaying increased usage of intronic polyA sites (lower part of the panel, 168 genes), merged from triplicates, in SCAF1-AID cells treated with (black line-SCAF1 depleted, control samples) and without (blue line-SCAF1 wildtype background) auxin.

To address whether SCAF1 directly associates with the affected genes, we analyzed previously published SCAF1 CUT&RUN data ^41^. Notably, we found that the genes affected by SCAF1 loss also showed a significantly higher degree of SCAF1 occupancy compared to input (Fig. 3e). Strikingly, we further observed that SCAF1 occupancy increased markedly towards the 3’ end of SCAF1 target genes identified through differential exon usage and with SCAF1-dependent polyadenylation site selection, placing SCAF1 at the site where we observed SCAF1-dependent transcription changes. These data further support the idea of a direct role for SCAF1 as a co-transcriptional regulator of PAS usage for a specific gene set in human cells.

### SCAF1 interaction with RNAPII is a prerequisite for its isoform switch phenotype

As SCAF1 contains an SRI domain, known from SETD2 to interact directly with the phosphorylated CTD of RNAPII ^42,43^, we next focused on characterization of this interaction to understand the basis for SCAF1’s interaction with RNAPII. Based on western blots using phosphorylation-specific RNAPII-CTD antibodies, we found that SCAF1 interacts with the RNAPII complexes enriched with Ser2, Ser5 and Ser7 phosphorylation (Fig. 1f, Supplementary Fig. 1c). However, since each RNAPII complex contains 52 heptad repeats it is not possible to conclude if this is due to a direct binding of either of these phosphorylation marks or simply because the marks are co-occurring on adjacent repeats. To investigate the specificity of SCAF1-SRI binding to the CTD repeats, phosphorylated CTD heptapeptide repeats (Fig. 4a) were synthesized and tested for binding to purified SCAF1-SRI (Supplementary Fig. 4a) using an *in vitro* fluorescence polarization assay. In accordance with the IP western blot results, we find that the SCAF1-SRI domain binds only to phosphorylated CTD peptides, with a strong preference for double phosphorylated Ser2-Ser5 RNAPII CTD peptides (Fig. 4b), which have recently been shown to typify elongating RNAPII ^44^. Thus, our data confirm a direct interaction between SCAF1-SRI and RNAPII CTD, with a binding preference for Ser2-Ser5 phosphorylated RNAPII CTD. Unlike previously reported, we did not observe any effect on RNAPII ubiquitination upon loss of SCAF1 (Supplementary Fig. 4b) nor did loss of SCAF1 lead to dramatic changes in the interactome of elongating RNAPII (Supplementary Fig. 4c and Supplementary Table S1) ^41^.

**Fig. 4:**
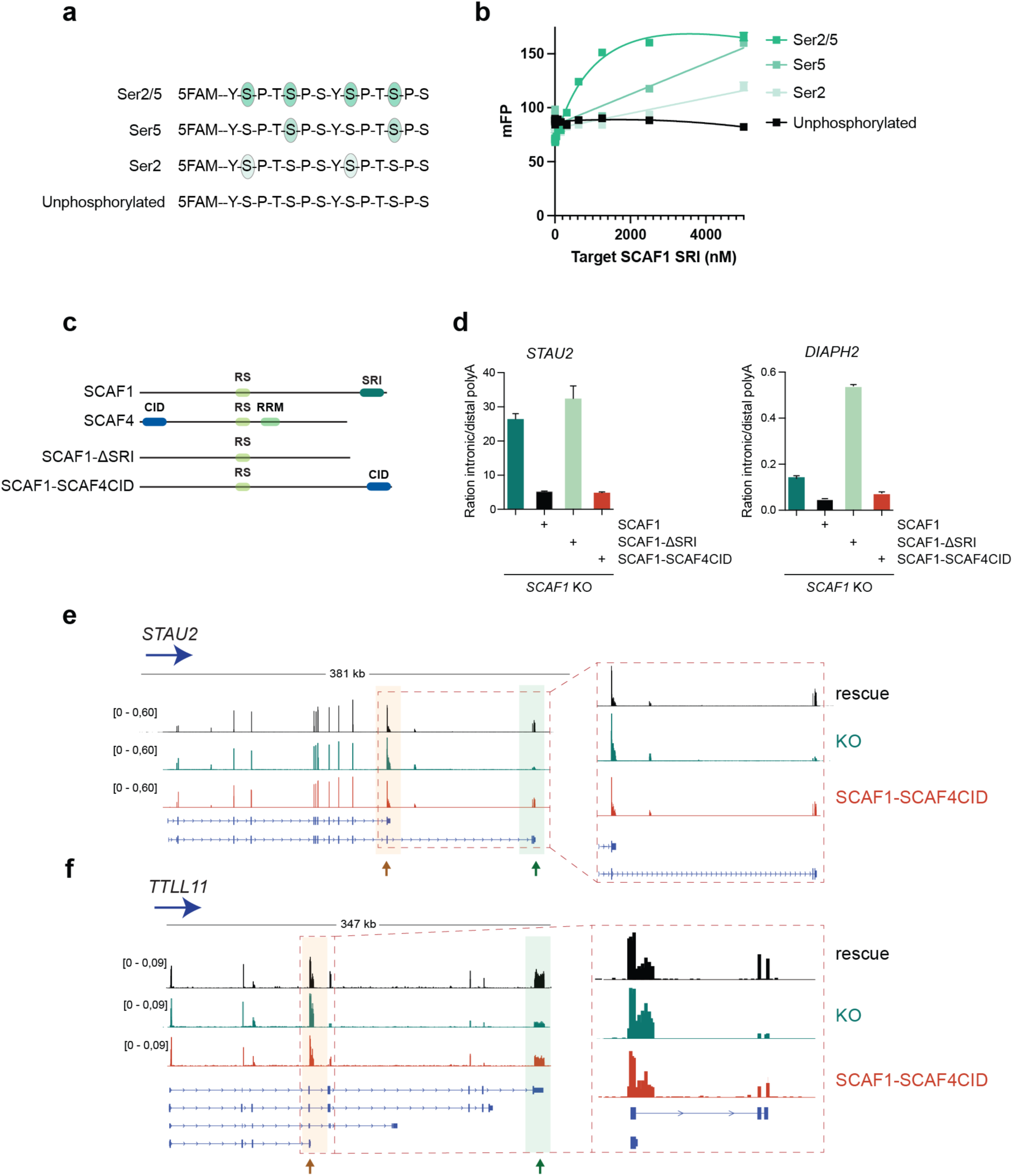
The SRI domain of SCAF1 interacts directly with the Ser2/Ser5 doubly phosphorylated RNAPII-CTD and is required for the transcriptional phenotype of SCAF1. **a** Amino acid sequence of the FAM-labeled RNAPII-CTD peptides used for the in vitro peptide binding assay. Differentially colored bubbles indicate the phosphorylated serines of the peptides. **b** Fluorescence polarization signal plotted as a function of wildtype SRI domain concentration for each probe (CTD peptide) included. The data were normalized to a blank, SRI lacking sample and the binding curves were generated from FP signal, as millipolarization units (mP), against the SRI protein concentration for each probe. Each experiment was conducted in duplicate. **c** Functional domains contained within SCAF1 and SCAF4 proteins, as well as SCAF1 variants used. Highlighted domains: **CID**: CTD interacting domain, **RS**: serine arginine rich region, **RRM**: RNA recognition motif, **SRI**: Set2 Rpb1 interacting domain). **d** Ratio of short to long mRNA isoform in the different SCAF1 variants (*SCAF1* KO, SCAF1 rescue, SCAF1-ΔSRI and SCAF1-SCAF4CID), validated by qPCR in two SCAF1 target gene examples. Primer-pairs used were designed to be specific for the short isoform 3’ UTR or the long isoform 3’ UTR. The ratio between products from these primers, normalized to *GAPDH* and a gene specific reference region (intron spanning primer pair common for both isoforms) is plotted as ratio of intronic to distal polyA. Error bars represent ±SD. **e, f** IGV tracks displaying mRNA-seq results from SCAF1 rescue, *SCAF1* KO and SCAF1-SCAF4CID cell lines. Orange and green arrows indicate the annotated sites of intronic and distal polyA sites respectively.

We next asked whether the SRI domain-mediated interaction between SCAF1 and RNAPII is required for the SCAF1 mediated polyadenylation site changes. To test this, we generated cell lines expressing SCAF1 lacking the SRI domain (SCAF1-ΔSRI). Interestingly, even though SCAF1 and SCAF4 both belong to the SCAF family, the domain interacting with RNAPII differs significantly structurally ^43,45,46^. We therefore also expressed SCAF1 with its SRI domain substituted by SCAF4’s CTD-interacting domain (SCAF1-SCAF4CID) (Fig. 4c). qPCR analysis of polyadenylation site usage in cell lines expressing the different SCAF1 variants revealed that SCAF1-ΔSRI is unable to reverse the usage of proximal polyadenylation site of the *SCAF1* KO background (Fig. 4d). On the other hand, substitution of SCAF1’s SRI domain with SCAF4’s CID domain was sufficient to restore the isoform switch phenotype observed in both in *STAU2* and *DIAPH2* (Fig. 4d). This was further confirmed with genome wide data, as mRNA-sequencing of cells expressing SCAF1-SCAF4CID perfectly clustered with the rescue SCAF1 samples (Supplementary Fig. 4f) and, as evident from single gene examples, the construct containing SCAF4’s CID fully rescues the phenotype of *SCAF1* KO cells for *STAU2* and another SCAF1-affected gene, *TTLL11* (Fig. 4e-f, Supplementary Fig. 4e). Moreover, the two rescue cell lines (SCAF1 and SCAF1-SCAF4CID) share a significant proportion of differentially expressed genes, as well as a significant overlap of genes found with alternative expression of exons, based on DEXseq analysis (Supplementary Fig. 4g). These data support the somewhat surprising conclusion that CTD-binding domains from different classes (ie SRI vs CID) can be interchangeable. They also further support that idea that the interaction between SCAF1 and RNAPII is necessary for supporting the gene-specific mRNA anti-terminator effects of SCAF1.

### SCAF1 dependent isoform switch is conserved in neuronal differentiation of mouse embryonic stem cells

Previous data showed that the ratio between long and short isoforms of transcripts can affect tissue specific gene expression and significantly impact cell fate and identity during differentiation and development ^6,47^. To address the role of SCAF1-dependent isoform switching in a model system permitting lineage-specific differentiation, we generated mESCs with endogenous SCAF1 harbouring an auxin-inducible degron tag (Fig. 5a). Degron-mediated depletion of SCAF1 was confirmed after 3 and 24 hours (hrs) in pluripotent mESCs cultured in metastable Serum-LIF media, that support self-renewal (Fig. 5b). 4sUmRNA-seq analysis of the newly synthesized full-length transcriptome showed that rapid degradation (3 hrs) of SCAF1 in pluripotent mESCs only resulted in relatively few differentially expressed genes and differential exon usage events (Supplementary Fig. 5a-d). To test the role of SCAF1 during the exit from pluripotency, we cultured the cells in N2B27 for lineage direction (either epiblast-like or spontaneous mixed differentiation), according to published experimental approaches ^48^. SCAF1 degron cells were cultured in different growth media 1) Naïve: 2iLiF containing media for maintained naïve pluripotent condition, 2) EpiLC: 2iLiF containing media supplemented with factors for exit from naïve pluripotency into epiblast-like cells and 3) Differentiation: 2iLiF removed media for different lineage differentiation, in the presence or absence of auxin (IAA) (Supplementary Fig. 5e) and were harvested after 2 days for mRNA-seq. Differential Gene Expression analysis (DGE) of the mRNA-seq data confirmed clustering based on culture conditions, as displayed in the overall heatmap (Supplementary Fig. 5f). GeneSet Enrichment Analysis (GSEA) of the genes differentially expressed in a SCAF1-dependent manner (+/- IAA) revealed that the vast majority of the affected pathways contained neuronal-related terms (Supplementary Fig. 5g).

**Fig. 5:**
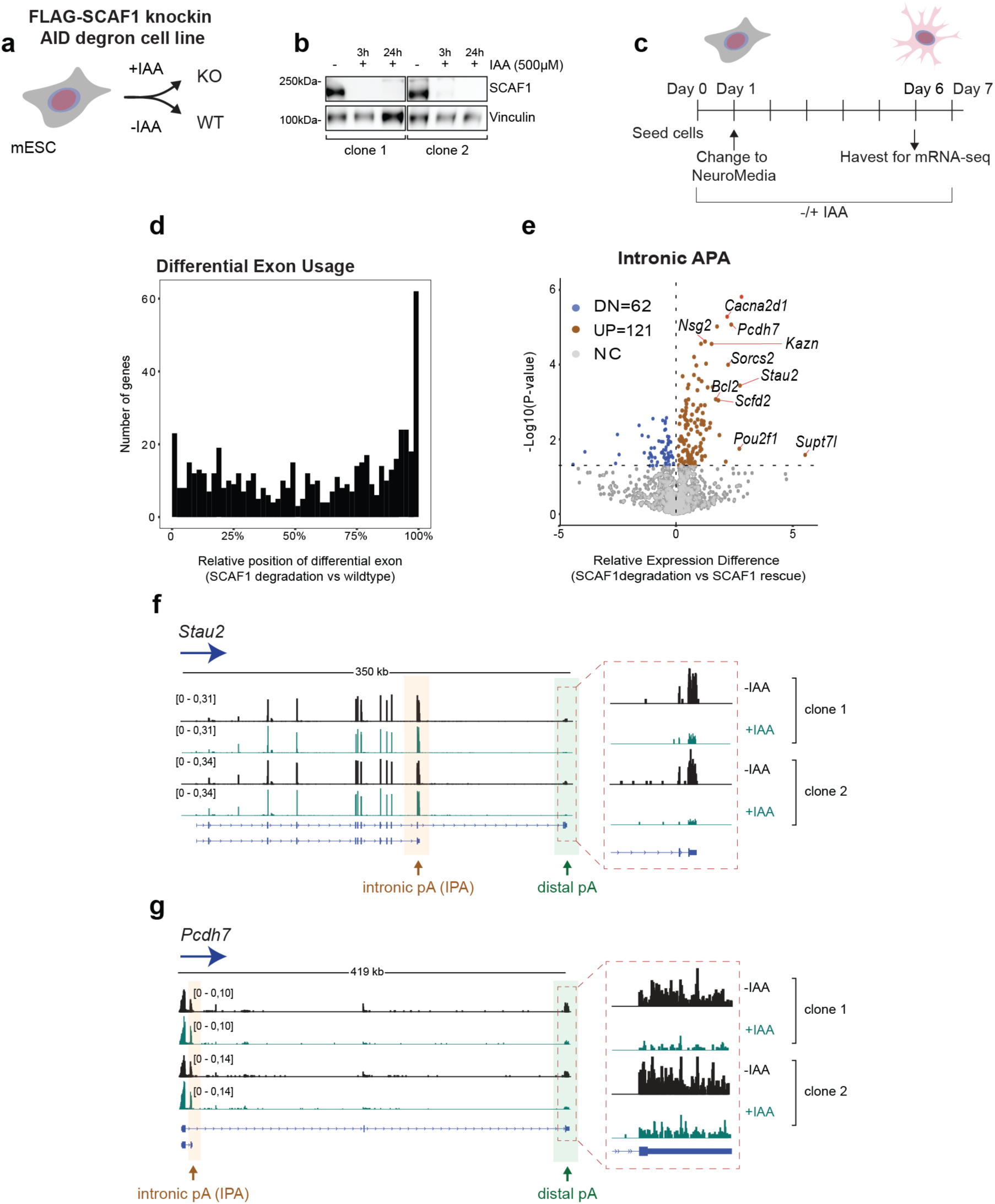
Loss of SCAF1 is conserved in the mouse cells and leads to upregulation of alternative intronic polyadenylation site usage. **a** Experimental set up for generation of degron cell lines in mESCs, resulting in *SCAF1* KO and wildtype conditions with the addition or not of Auxin (IAA) respectively. **b** Western blot verifying SCAF1 degradation in 3 and 24 hours (hrs) after addition of 500 μM IAA in two clones used. Vinculin is used as loading control. **c** Graphical outline of the neuronal differentiation protocol. On day 0, two individual SCAF1 degron cell lines are seeded in the presence or not of IAA and incubated for 24 hrs, prior to switch to neuronal differentiation media (NeuroMedia), in which they are cultured for the following 5 days. On day 6, cells are harvested for mRNA-seq or used for imaging to confirm successful differentiation. **d** Barplot displaying the relative position of differential exon usage, upon degradation of SCAF1 in differentiated mESCs (DEX-seq analysis). **e** Volcano plot, generated after APAlyzer analysis, showing differentially used intronic APA sites in SCAF1 depletion, compared to wildtype conditions. Data were merged from two clones, sequenced in triplicates per *SCAF1* genotype background. Colored dots represent transcripts with significantly (p-value <0.05) increased (dark red) or decreased (blue) usage of the annotated intronic APA sites. **f, g** IGV tracks displaying mRNA-seq results from two SCAF1 degron clones tested with wildtype (-IAA) and SCAF1 degraded (+IAA) backgrounds. Orange and green arrows indicate the annotated sites of intronic and distal polyadenylation (polyA) sites respectively. Gene examples were also identified through APA analysis.

Since mRNA isoform changes and alternative polyadenylation site usage has frequently been observed during neuronal differentiation ^5,49–51^, we investigated whether SCAF1 affects gene expression of neuronal genes, and if so, whether this is directly related to changes in mRNA isoform expression via polyadenylation site choice. To address this, we drove differentiation of naïve mESCs towards the neuronal lineage, culturing them in Neuromedia (Advanced DMEM/F12 supplemented with NEA) and detected genome-wide changes through transcriptome profiling (Fig. 5c). Comparison of samples with and without SCAF1 expression revealed that, similar to our observation in human cells, SCAF1 dependent mRNA isoform switching was also observed in differentiated mouse cells (Fig. 5d-g). Again, upon loss of SCAF1, we detect a set of genes with differentially affected exons, particularly the last ones (Fig. 5d), and with increased usage of intronic polyadenylation sites (Fig. 5e, Supplementary Fig. 5h). This was also evident from single gene examples (Fig. 5f-g, Supplementary Fig. 6d). Interestingly, a subset of 25 genes regulated by SCAF1 in HEK293 cells was also SCAF1 regulated in differentiated mouse cells, points to evolutionary conservation of SCAF1 function (Supplementary Fig. 5i). Together both data from human and mouse loss of function SCAF1 models suggest a role in regulation of mRNA isoform expression, through regulation of PAS selection, particularly for PAS located towards the end of genes.

### SCAF1 affects neuronal gene expression and is required for neuronal differentiation

We next investigated whether the effects of SCAF1 on polyA site selection has an impact on neuronal differentiation. Indeed, loss of SCAF1 protein led to a differential expression of hundreds of genes, with a profound downregulation of genes in pathways related to neuronal activity and function (Fig. 6a). Importantly, transcription factors critical for neuronal lineage commitment, such as *Ascl1*, were also suppressed upon SCAF1 degradation, indicating a broad impairment of neurogenic programming (Fig. 6b-c)^52^. In contrast, the genes upregulated upon SCAF1 degradation showed some enrichment for immune response related terms (Supplementary Fig. 6a).

**Fig. 6:**
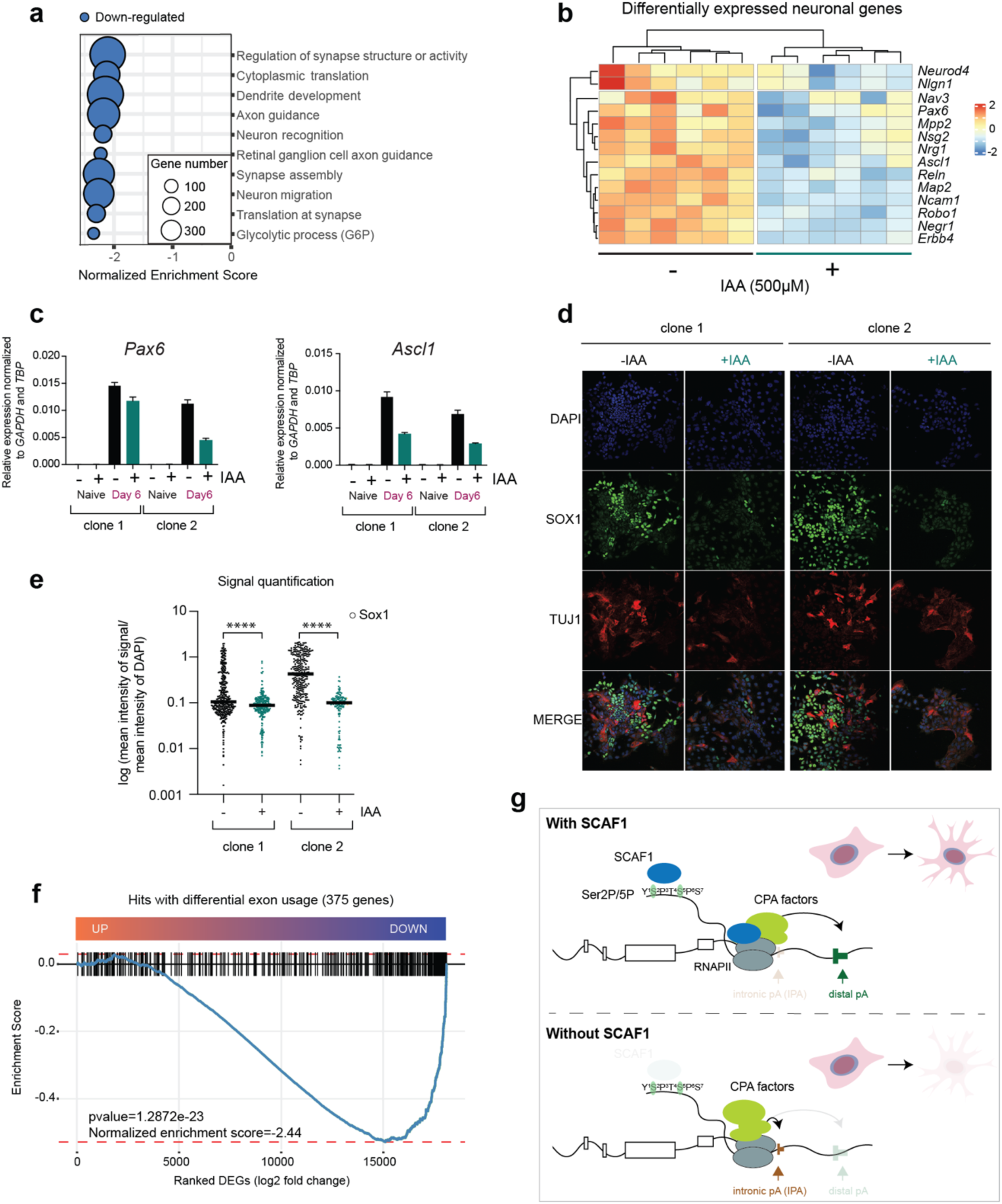
SCAF1 regulates polyadenylation site selection of neuronal genes and drives neuronal differentiation. **a** Bubble plot from gene set enrichment analysis of the expression changes (SCAF1 degradation vs wildtype). The top 10 most significantly downregulated pathways are shown, with normalized enrichment scores (padj < 0.05) and bubble sizes scaled accordingly to number of downregulated genes within individual categories. **b** Heatmap depicting differentially expressed neuronal genes identified from mRNA-seq data of neuronal differentiated samples harvested in triplicates from two SCAF1 degron cell lines. Difference in expression is illustrated based on log2FC of selected neuronal genes between SCAF1 expressing (-IAA) and SCAF1 degraded (+IAA) samples at day 6 of neuronal differentiation. **c** qPCR validation of the expression pattern of two neuronal genes in the two SCAF1 degron clones. Naïve/pluripotent samples were used as control for expression induction of the genes during differentiation. *Gapdh* and *Tbp* were used as internal control. Samples were run in quadruplicates and error bars represent ±SD. **d** Immunofluorescence (IF) images captured at 40x magnification, using the nuclear, neuronal stem cell marker SOX1 and the marker for early neurons, TUJ1, in the two SCAF1 degron clones. The experiment was performed twice. **e** Quantification of nuclear SOX1 signal from the IF experiment described in (**d**). Violin plots of the different clones presenting the log10 mean intensity of SOX1 signal, normalized to background DAPI intensity. Statistical significance is indicated with asterisks (mann-whitney unpaired t-test with pvalue<0.0001, ****). **f** Pre-ranked gene set enrichment analysis of mRNA-seq (x-axis), from the most upregulated to the most downregulated genes identified from DESeq2 analysis. Blue lines show the trend of the genes with either differential exon usage, according to differential expressed genes. Statistical significance (p-value) and normalized enrichment score are indicated. **g** Model for SCAF1 action in co-transcriptional processing and how it affects neuronal differentiation. SCAF1 associates with RNAPII, that is accessorized with active elongation and 3’ end processing factors, through the interaction of its SRI domain with Ser2/Ser5 bi-phosphorylated RNAPII CTD and regulates the usage of polyA sites for a subset of genes. The role of SCAF1 is required particularly during neuronal differentiation, as it secures tight control of the precise synthesis of neuronal mRNA transcripts. Even though lack of SCAF1 leads to shift towards usage of proximal polyA sites of more than 100 genes, knockout of SCAF1 in HEK293 cells is not accompanied with major morphological or growth phenotypes; however, the effects of SCAF1 loss is obvious during neuronal differentiation, with a clear cellular phenotype. In neuronal differentiating mESCs, SCAF1 affects neuronal gene expression by promoting the usage of polyA site; upon SCAF1 degradation, a subset of neuronal genes expresses truncated mRNA transcripts, resulting to downregulation of their expression contributing synergistically to neuronal differentiation defects.

Neuronal deregulation was further validated by immunostaining for TUJ1, a pan-neuronal marker of post-mitotic neurons, together with SOX1, a marker of neuroectoderm and neural progenitors. Upon SCAF1 loss, we observed a significantly reduced proportion of SOX1-positive cells, suggesting that SCAF1 depletion perturbs the maintenance of the neural progenitor pool and leads to aberrant neuronal differentiation (Fig. 6d-e, Supplementary Fig. 6c). Exit from pluripotency in SCAF1 degron cell lines was validated with qPCR analysis, displaying reduced to minimal expression of naïve markers (*Nanog* and *Pou51*), confirming that cells lacking SCAF1 are able to exit naïve pluripotency. However, despite successful exit from the pluripotent state, SCAF1-depleted cells failed to efficiently activate neuronal-specific gene expression, indicating a block in the execution of the neurogenic program (Supplementary Fig. 6b).

Interestingly, SCAF1 target genes, detected both by DEXseq and IPA, were significantly enriched among the downregulated genes, as evident from the negative enrichment score upon Gene Set Enrichment Analysis (GSEA). (Fig. 6f, Supplementary Fig. 6d). As shown previously, the expression of neuronal related genes is highly downregulated. Indeed, in several cases, it was evident that the 3’ end of neuronal genes was specifically affected, with elevated usage of proximal polyadenylation sites, likely explaining the inability of mouse cells to commit to neuronal differentiation due to deregulations in SCAF1-dependent co-transcriptional APAs (Fig. 5f-g, Supplementary Fig. 6e). Among the SCAF1 targets was *Pcdh7*, which encodes Protocadherin 7, highly expressed in the brain and heart ^53^ and shown to drive neuronal stem cell differentiation ^54^, and *Kazn*, which expresses the Kazrin protein, critical for brain development in *Xenopus* ^55^. Moreover, a common SCAF1 target identified in both human and mouse cell lines -*Stau2*-encodes the RNA binding protein Staufen 2 (STAU2) protein, crucial for dendritic spine morphogenesis ^56^. Therefore, SCAF1 driven co-transcriptional APA regulation of neuronal genes leads to downregulation of their expression, an effect that likely leads to defect in neuronal differentiation.

Together, these findings show that SCAF1 controls usage of the polyadenylation sites in the target transcripts, a role that is crucial in the regulation of neuronal gene expression and neuronal differentiation (Fig. 6g). Loss of SCAF1 thus leads to a shift towards usage of proximal polyA sites in the target genes and severely affects neuronal differentiation, a process that is highly dependent on the controlled processing of mRNA isoform 3’ends.

## Discussion

Although SCAF proteins were initially identified in the mid-90’ as RNAPII CTD interactors ^29^, their function has, for most parts, remained elusive. Here, we find that SCAF1 acts as a regulator of polyadenylation site usage, similarly to SCAF4 and SCAF8, which we have previously shown to act as mRNA anti-terminator by suppressing use of proximal PAS ^21^. While both SCAF4 and SCAF8 target early intronic PAS, often located in the beginning of genes, SCAF1 most often affects PAS usage in exons towards the 3’end of genes, pointing to a distinct mechanism from SCAF4/SCAF8. Differences in action perhaps are not surprising, given the diversity within the SCAF family. Although they all interact directly with the phosphorylated RNAPII CTD, they do not share overall sequence similarities, with exception of SCAF4 and SCAF8 which arose from a gene duplication ^21^. While SCAF4 and SCAF8 contain a CTD interacting domain (CID) also found in PCF11, Seb1, Nrd1, and Rtt103 ^46,57^, SCAF1 contains the structurally distinct Set2 Rpb1 Interacting domain (SRI) also found in SETD2, RECQL5, PHRF1 and SCAF11 ^43^. Due to this difference in the CTD-interacting domain between SCAF1 and SCAF4/SCAF8, we would have expected that this would impact the specificity of the RNAPII interaction, making the CID and SRI containing SCAF proteins somewhat distinct in terms of their RNAPII binding. Despite this, SCAF1, similarly to SCAF4/SCAF8 has a high affinity towards the doubly Ser2P/Ser5P RNAPII CTD, which also explains why we find SCAF1 bound to the same RNAPII complexes as SCAF4 and SCAF8. This suggest that the same ‘kind’ of RNAPII CTD-binding has evolved from different directions, through structurally distinct domains. In this regard, it is interesting to note that the doubly Ser2P/Ser5P RNAPII CTD mark is predominantly found in higher eukaryotes, while almost absent in yeast ^58^. It has been speculated that this is related to the increased gene length in higher eukaryotes, thus making the Ser2P/Ser5P specific to the long elongation phase in higher eukaryotes ^58^. This again hints that the SCAF family has evolved to fine-tune the transcriptional output in a manner specifically required in higher eukaryotes, both in regard to their RNAPII interaction and mode of regulation. Our finding that SCAF1 containing a SCAF4-CID domain in place of its own SRI domain can rescue the SCAF1-dependent transcriptional phenotype, suggests two key points: First, that even despite their difference between the CID and SRI domains, they serve the same functional purpose, which is recruitment of the SCAF proteins to RNAPII. Second, that the functional differences in target site selection between SCAF1 and SCAF4/SCAF8 must originate from the remaining part of the proteins. Beside the SRI and a short arginine-serine rich domain, SCAF1 does not contain any annotated domains, and structural predictions indicate no presence of ordered regions. Instead, SCAF1 harbors several low complexity regions that might be involved in protein-protein interactions, potentially through liquid-liquid phase separation. We hypothesize that such interactions could be achieved with other CPA factors, as recently shown for the CPA component RBBP6 ^59–61^. Mechanistically, we show that SCAF1 interacts with both RNAPII bound by elongation complexes, such as PAF1 complex, and RNAPII complexes enriched for binding to numerous CPA factors, including, CPSF, CFI, CFII and CStF. Such results suggest that SCAF1 is indeed associating with RNAPII complexes as they transition from active elongation into 3’end mRNA cleavage and termination. In agreement with this, we find SCAF1 binding enriched towards the 3’end of genes, for which we have identified SCAF1-dependent PAS changes, again fitting with a model where SCAF1 marks a subset of RNAPII complexes competent for 3’end mRNA cleavage. While SCAF1 was recently identified in a screen to affect global transcription levels ^41^, we do not observe any changes in overall RNAPII transcription levels, either based on spike-in normalized nascent transcriptomics or EU labelling upon SCAF1 depletion. Additionally, we do not find any evidence of drastic changes to the RNAPII interactome, upon loss of SCAF1. These observations are very much in line with previous findings for *SCAF4* and *SCAF8* KOs, that similarly displayed nascent transcription changes restricted to a defined subset of genes, with changes occurring downstream of SCAF-dependent mRNA cleavage events ^21^.

While SCAF4 and SCAF8 suppress usage of early intronic PAS in several hundred genes, the number of SCAF1-affected genes is more selective. This difference between SCAF1 and SCAF4/SCAF8 is further highlighted by the phenotypic differences observed in the two setups: loss of SCAF1 in HEK293 cells does not affect cellular growth, while double knockout of *SCAF4* and *SCAF8* is lethal ^21^. While we identify overall 265 and 183 SCAF1-regulated APA changes in HEK293 and mouse cells respectively, considering that most SCAF1-regulated genes are not expressed in both HEK293 and mouse cells, this relatively small but statistically significant overlap of 25 genes nevertheless suggests a conserved role for SCAF1 in regulating the polyadenylation site choice of target genes also in mammals.

Importantly, we report that SCAF1 is functionally required during finely tuned developmental programs, as during neuronal differentiation, that relies on highly coordinated temporal expression of tissue-specific mRNA isoforms. Lack of SCAF1 during neuronal differentiation of mESCs cells leads to mis-regulation of numerous genes required for neurogenic program and, consequently, cells lacking SCAF1 fail to differentiate efficiently. In line with its functional role during neuronal differentiation, expression levels of SCAF1 are the highest in the fetal brain ^62^, further supporting its importance in regulating the precise expression of neuronal genes.

Extensive work has focused on mRNA isoform expression, specifically during neuronal differentiation ^5,51,63,64^. However, factors regulating PAS selection remain poorly understood. The best studied example include the neuronal RNA-binding protein (nRBP) ELAV/Hu family proteins being involved in determining neuronal identity via regulation of APA within the 3’UTR ^65,66^. In addition, SPOC domain containing PHF3 and DIDO1 have recently been described to regulate a specific subset of genes with an importantly role during neuronal differentiation ^67–69^. In the case of PHF3, this is achieved through regulation of RNAPII elongation rate and mRNA stability ^67^, neither of which we found to change in cells lacking SCAF1 (data not shown), suggesting a different mode-of-action between SCAF proteins and SPOC domain factors, such as PHF3. Our findings highlight SCAF1 as new mode of APA regulation of neuronal genes, carried out co-transcriptionally through specific recruitment to RNAPII subcomplexes towards the end of genes, impacting last exon choice. In fact, distal last exon control is reported to promote localization of corresponding transcripts to neurites ^24^, Therefore one possibility is that SCAF1 mediated APA regulation serves to ensure proper processing of neuronal mRNAs that need to translocate.

Unlike SCAF4 and SCAF8 that evolved from the yeast homolog Nrd1 and Seb1, SCAF1 is only present in vertebrates, supporting its function in regulating more specialized mRNA processing pathways, such as PAS selection in specific cell types, like neurons. This suggests that SCAF1 has evolved to serve a crucial role, in regulation of mRNA isoforms by bridging RNAPII CTD recognition with co-transcriptional mRNA processing, something uniquely required in higher eukaryotes and particularly during development. Although *SCAF1* expression is the highest in the brain, it is expressed across all tissue types, leaving the possibility that SCAF1 is also involved in the regulation of mRNA isoform expression during other developmental programs. Recent work on the CFI complex subunit, CPSF6, shows a clear separation of polyadenylation effects, in which loss of CPSF6 in the brain versus heart leads to increased events of alternative last exons and alternative distal 3’UTR respectively ^47^. Thus, it would be interesting to test the effect of SCAF1 loss in other tissues, like the heart and whether SCAF1 follows a different, tissue specific mRNA processing regulation. Interestingly, SCAF1 has been suggested as a pancreatic tumor suppressor together with USP15 ^31^. Curiously this observation was linked to upregulation of truncated USP15 protein expression in a SCAF1-dependent manner ^31^. This is in strong agreement with the results from HEK293 cells showing that SCAF1-mediated PAS usage in the case of USP15 leads to the production of a truncated cytoplasmic form of USP15 and downregulation of nuclear full-length USP15. Together, this further validates the functional importance of SCAF1-mediated PAS selection and suggests that SCAF1-mediated regulation could play importantly functional role across multiple tissues and, as reported, drive changes in cancer genes.

In summary, our work highlights the surprising finding that SCAF1, although structurally and sequence-wise distinct from SCAF4 and SCAF8, share a functional commonality, which is the regulation of polyadenylation site usage. This function is shared with the *Schizosaccharomyces pombe* homolog of SCAF4 and SCAF8, Seb1 ^70,71^, but distinct from the *Saccharomyces cerevisiae* Nrd1 protein, which, despite containing the same functional domains as SCAF4, SCAF8 and Seb1, has an entirely different functional role as part of the Nrd1-Nab3-Sen1 complex in transcriptional termination of cryptic transcript ^72,73^. While the yeast CID-containing proteins Nrd1 and Seb1 associate with RNAPII Ser5P or Ser2P, respectively ^71,73^, the so far characterized human SCAFs (SCAF1, SCAF4 and SCAF8) all share a preference for the double phosphorylated Ser2P/Ser5P elongating RNAPII and regulate PAS usage either early in the gene as observed for SCAF4 and SCAF8 ^21^ or towards the 3’end as described here for SCAF1. Curiously, the RNAPII CTD interaction is achieved either though the CID (in case of SCAF4 and SCAF8) or an SRI domain (in case of SCAF1), which serves and interchangeable function; recruitment to RNAPII, suggesting that the SCAFs have evolved independently towards common functionality in higher eukaryotes, representing a new class of proteins modulating PAS usage.

## Methods

### Cell lines and culture conditions

Flp-In T-REx HEK293 cells (Thermo Fisher Scientific, R78007, human embryonic kidney epithelial, female) and Flp-In T-REx U2OS (female, gift from Michael Lisby group) were cultured in high glucose DMEM (Thermo Fisher Scientific, 11965118) supplemented with 10% (v/v) FBS, 2 mM L-glutamine, 100 U/mL penicillin, 100 μg/mL streptomycin, and 15 μg/mL blasticidin at 37°C with 5% CO_2_.

Naive mouse embryonic stem cells (mESCs) were cultured in fibronectin-coated plates (Merck - 1/60) N2B27 basal culture medium (in 1:1 DMEM/F12 medium - Life Technologies and Neurobasal medium - Gibco, N2 supplement 1/200 - Gibco, B-27 Serum-Free Supplement 1/100 - Gibco, 2mM L-alanyl-L-glutamine - GlutaMax, Gibco, 0.1 mM 2-mercaptoethanol - Sigma) supplemented with 2iLiF (3 μM of CHIR99021, 1 µM of PD0325901, and 1,000 U mL−1 of LIF produced in-house).

### Plasmid construction and generation of stable cell lines

*SCAF1* transcript was gene synthesized through GenScript and cloned in pENTR4 backbone vectors (pENTR4_SCAF1). Fragment corresponding to SRI domain of SCAF1 was removed also through GenScript and cloned in pENTR4 vectors (pENTR4_SCAF1-ΔSRI). SCAF1-SCAF4CID construct was created with Gibson assembly, using BmgBI/NotI digested pENTR4_SCAF1 backbone and swapping the SRI corresponding fragment to the amplified from CID fragment from pFRT/TO/SCAF4-FLAG generated at ^21^. All *SCAF1* constructs were recombined with pDEST/TO/FLAGHA or pDEST/TO/GFP with epitope tag inserted at the N-terminus of the SCAF1 transcript using Gateway LR Clonase II Enzyme mix according to the manufacturer’s protocol (Thermo Fisher Scientific, 11791020). *SCAF1* knockout was achieved through CRISPR Cas9 technology, with gRNA targeting the seventh exon in *SCAF1* gene. The designed gRNA was cloned into BbsI digested CRISPR Cas9-GFP tagged plasmid pX458 (Addgene plasmid #48138). HEK293 cells were transfected with pX458 and 48 hrs later, GFP-positive cells were single cell sorted using FACS Melody sorter and were incubated for 12 days. Genomic DNA from single colonies was extracted, exon 7 was amplified with primers designed to generate product of 700 bp that included the BAM sequence. Samples were Sanger sequenced, together with wildtype control DNA, and analyzed for potential inserts/deletions using online tool TIDE (Tracking of Indels by Decomposition). *SCAF1* KOs were confirmed with western blot using SCAF1 antibody. Stable, epitope expressing SCAF1 cell lines were generated by integration of respective constructs into the FRT locus of HEK293 Flp-In TREx or U2OS Flp-In TREx cells, as described at ^21^. Briefly, constructs (FLAGHA/TO-SCAF1, FLAGHA/TO-SCAF1-ΔSRI, GFP/TO-SCAF1-SCAF4CID) were co-transfected with pOG44 Flp-recombinase expression vector (Thermo Fisher Scientific, V600520) in a 9:1 ratio. 24 hrs after transfection, cells were expanded to 15 cm dishes and after another 24 hrs the cell culture media was replenished to media supplemented with 100 mg/mL hygromycin and 15 mg/mL blasticidin. 12 days after selection, single clones were picked and expanded. Expression of tagged proteins was induced with the addition of doxycycline for 48 hrs (Clontech, 8634-1, 0.1 µg/µL final concentration). Western blot verification of the different cell lines was performed using the both the SCAF1 and HA or GFP antibodies.

For generation of SCAF1-mAID-FLAG degron cell system (SCAF1 degron) we used E14Tg2a (129/Ola,XY) wild-type mESCs that were stably transfected with the OsTir1 gene at the Igs7(Tigre) locus, under the constitutive CAG promoter. Genetic engineering of the *Scaf1* locus introduced the degron tag at the C-terminus of *Scaf1* and was performed by nucleofection of cells using the P3 Primary Cell 4D-Nucleofector™ X Kit (Amaxa - Lonza). SCAF1 degron clones were verified with western blot using SCAF1 antibody, comparing SCAF1 protein signal between wildtype and SCAF1 3 hrs degradation (with addition of 500µM IAA), followed by karyotyping to assess correct chromosome number and integrity.

### EpiLC and Neuronal induction conditions

All differentiations were performed as described in the following ^48,74,75^. For exit from pluripotency and spontaneous differentiation, cells were cultured for two days in N2B27 basal culture medium, without 2iLiF supplementation. For EpiLC induction, cells were cultured for two days in N2B27 basal culture medium supplemented with bFGF (12 ng/mL) and Activin A (20 ng/mL) and 1% KSR. Medium (+/- IAA) was changed on a daily basis. Neuronal differentiation was performed in laminin-coated plates for 6 days upon 2iLiF withdrawal, in N2B27 neuro medium (referred as Neuromedia): mix of 1:1 Advanced DMEM/F12 (Thermo - 12634010) and Neurobasal medium (Thermo - 21103049), supplemented with 1x GlutaMax (Thermo - 35050061), 1x NEA (Thermo - 11140035), 0.5x N2 supplement (Thermo - 17502048), 0.5x B27 supplement (17504044) and 0.1 mM 2-mercaptoethanol. Cells were seeded in the corresponding culture conditions, with the addition or not of 500μM IAA, and culture media was replenished, for the number of days indicated at the treatment.

### Cell fractionation

HEK293 wildtype cells expressing FLAGHA-SCAF1, upon 0.1 µg/µL doxycycline induction, were seeded in a 15 cm dish. Cells were scraped with PBS and centrifuged at 300 g for 4 min at room temperature. Cell pellets were incubated in 4 pellet volumes of soluble extraction (SE) buffer, (20 mM HEPES-KOH pH 7.5, 150 mM potassium acetate, 1.5 mM MgCl₂, 0.5% (v/v) NP-40, 10% (v/v) glycerol, protease and phosphatase inhibitors) on ice for 20 min and then spun down at 21000 g for 20 min at 4°. Supernatant was collected as soluble (**S**) fraction. Nuclear pellets were resuspended in 2 pellet volumes of chromatin dilution buffer (20 mM HEPES-KOH pH 7.5, 150 mM NaCl, 1.5 mM MgCl₂, 0.05% (v/v) NP-40, 10% (v/v) glycerol, protease and phosphatase inhibitors) and incubated for 1 hr rotating at 4°. Low chromatin (**LC**) fractions were isolated from supernatant, after centrifugation at 21000 g for 20 min at 4°. Proteins tightly bound to chromatin were extracted from resuspended cell pellets in the following steps: resuspension in 0.6 volumes of high salt buffer (20 mM HEPES-KOH pH 7.5, 500 mM NaCl, 1.5 mM MgCl₂, 0.05% (v/v) NP-40, 10% (v/v) glycerol, 0.1% (v/v) DNARASE®, protease and phosphatase inhibitors), incubated for 1 hr rotating at 4°, followed by the addition of 1.4 volumes of dilution buffer (20 mM HEPES-KOH pH 7.5, 1.5 mM MgCl₂, 0.05% (v/v) NP-40, 10% (v/v) glycerol, 0.1% (v/v) DNARASE®, protease and phosphatase inhibitors). Samples were centrifuged at 21000 g for 20 min at 4°C and supernatant was collected as high chromatin (**HC**) bound protein extract. The remaining proteins were extracted with Urea buffer (8 M Urea in 20 mM HEPES-KOH pH 7.5) incubated at room temperature for 15 min, spun down and supernatant collected as Urea (**U**) fraction.

### Microscopy

For human cells, GFP-SCAF1 expressing U2OS cells were seeded in the presence of 0.1 µg/µL doxycycline in 6-well plates with added coverslips, pre-coated in poly-lysine (Gibco). Once cells reached 60% confluency, media was replaced with 4% (v/v) formaldehyde in PBS. Cells were fixed at room temperature for 15 min, followed by mounting onto slides using VECTASHIELD Antifade Mounting Medium containing DAPI (Vector Laboratories, H-1200) and visualization using an inverted LSM 780 laser scanning confocal microscope (Zeiss).

For mouse cell lines, 500 cells were seeded in laminin-coated (10 µg/mL in PBS) 8-well μ-Slide Ibidi dishes (Ramcon, 80826) using 2iLiF containing media for keeping cells in naïve state. After 6 days in neuronal differentiation media, cells were fixed with 4% formaldehyde in PBS for 10 min at room temperature and washed three times with PBS. Then we permeabilized with 0.25% Triton X-100 for 10 min at room temperature, followed by three washes with PBS. Samples were blocked with blocking buffer (1% BSA, 0.1% Tween20, 10% Normal Donkey Serum in PBS, 10X PBS and H2O) for 3 hrs at room temperature and were incubated overnight at 4°C with primary antibodies diluted in blocking buffer. Samples were then washed three times in PBS-T (0.1% (v/v) Tween20 in PBS) and incubated with the secondary antibodies diluted in blocking buffer for 3 hrs at room temperature. DAPI was added after three washes with PBS-T and the samples were imaged on a Leica Stellaris 8 confocal microscope.

### Growth rate analysis

10000 cells per condition (in the presence or absence of 0.1 µg/µL doxycycline) were seeded with (tetracycline-free) FBS containing media in triplicates in a 96-well plate. The growth was monitored using an IncuCyte S3 Live-Cell Analysis System (Sartorius) and images of live cells were captured every 3 hrs for up to 5 days. Cell confluency (%) was measured with the IncuCyte S3 image analysis software.

### Western blotting

Cells were seeded in same numbers and were lysed for protein extracts with Urea buffer (8 M Urea in 20 mM HEPES-KOH pH 7.5) at room temperature for 15 min. Specifically for SCAF1 blotting samples, a prior step of soluble fraction removal (as described at cell fractionation section) was performed, to enhance the SCAF1 antibody specificity. Proteins were separated in either 6% or 15% homemade acrylamide gels and transferred to nitrocellulose membranes (GE Healthcare Life Sciences, 10600002). Blocking was performed in 5% (w/v) skimmed milk in TBS-T (TBS, 0.1% (v/v) Tween20) for 1 hr at room temperature followed by primary antibody (in 5% (w/v) skimmed milk in TBS-T) overnight incubation at 4°C. Vinculin, LaminA and Histone H3 served as loading controls. Membranes were washed several times in TBS-T, incubated with HRP-conjugated secondary antibody diluted in 5% (w/v) skimmed milk in TBS-T and visualized using Clarity and Clarity Max ECL Western Blotting Substrates (Biorad, 1705061 or 1705062).

### FLAGHA-SCAF1 immunoprecipitations

FLAGHA-SCAF1 cells in HEK293 background were seeded in 5×15 cm dishes per replicate (triplicates) and expression was induced with the addition of 0.1 µg/µL of doxycycline, 48 hrs prior to harvesting. HEK293 wildtype cells were included with the same seeding conditions and were used as negative control for the immunoprecipitation. Cells were harvested and chromatin fractionation was performed, as described on cell fractionation section, to enrich for SCAF1 abundance. **LC** and **HC** fractions from each condition were pooled, with input samples collected, followed by immunoprecipitation using monoclonal FLAG antibody (Sigma-Aldrich, F1804) bound to protein G dynabeads (Thermo Fisher, 10003D). For conjugating the FLAG antibody to dynabeads: 80 µL of soluble beads were used per replicate and were washed 3 times with binding and washing buffer (5 mM Tris-HCl pH 7.5, 0.5 mM EDTA, 1M NaCl, 0.1% (v/v) Tween20) and 3 times with PBS. 5 µg of FLAG antibody in 1 mL PBS were added to each of the beads and incubated at room temperature for 45 min. After incubation, beads were washed 3 times with 0.1% (v/v) Tween20 in PBS.

Chromatin extracts were added to the beads and incubated for 3 hrs at 4° C, rotating. Beads were washed 3 times with high KCL buffer (20 mM HEPES-KOH pH 7.9, 300 mM KCl, 0.2 mM EDTA, 20% (v/v) glycerol, 0.1% (v/v) Tween20) and 3 time with low KCL buffer (20 mM HEPES-KOH pH 7.9, 100 mM KCl, 0.2 mM EDTA, 20% (v/v) glycerol, 0.1% (v/v) Tween20). For SDS-PAGE analysis, immunoprecipitants were eluted from the beads by boiling them with 100 µL 2xSDS sample buffer and isolating the supernatant.

### RNAPII immunoprecipitation

HEK293 doxycycline inducible *SCAF1* KO clone was seeded in 10×15 cm dishes per replicate (n=3, 30 dishes total), in (tetracycline-free) FBS containing media, with half of the plates having doxycycline (0.1 µg/µL) added. Cells harvesting and chromatin fraction extraction and pooling procedure is the same as the one described at FLAGHA-SCAF1 immunoprecipitations section. Immunoprecipitation of phosphorylated RPB1 was performed using 2 µg of 4H8 antibody conjugated to protein G dynabeads (Thermo Fisher, 10003D). Antibody conjugation to beads and the rest of the immunoprecipitation procedure followed is described in FLAGHA-SCAF1 immunoprecipitations section.

### Mass spectrometry

Immunoprecipitation (IP) samples for proteomic analysis were performed in triplicate, with all samples prepared in parallel starting from cell seeding. Frozen, washed beads were thawed and incubated for 30 minutes with Elution Buffer 1 (2 M urea, 50 mM Tris-HCl pH 7.5, 2 mM DTT, 20 mg/ml trypsin), followed by a 5-minute incubation with Elution Buffer 2 (2 M urea, 50 mM Tris-HCl pH 7.5, 10 mM chloroacetamide). The resulting eluates were combined and incubated overnight at room temperature. Tryptic peptides were acidified to 1% TFA and loaded onto Evotips (Evosep). Peptide separation was carried out using 15 cm × 150 µm ID columns packed with 1.9 µm C18 beads (Pepsep) on an Evosep ONE HPLC system using the ‘30 samples per day’ workflow. Peptides were introduced via a CaptiveSpray source with a 10 mm emitter into a timsTOF Pro mass spectrometer (Bruker) operating in PASEF mode.

### SCAF1 knockdown (siRNA)

U2OS and HEK293 wildtype cells were seeded in antibiotic-free media in 6 well plates at low density (2.5×10^5^ cells) and incubated at 37°C overnight. Knockdown was performed with the addition of diluted in serum-free media of 20 nM siRNA (control or SCAF1) and 1:1000 Lipofectamine RNAiMAX (Thermo). Cells were incubated in the siRNA mix for 48 hrs and subsequently used for EU labelling, to quantify nascent transcription levels.

### EU labelling

Cells were seeded in pre-coated with poly-lysine (Gibco) Greiner CELLSTAR (Sigma-Aldrich, M0562-32EA), at a density of 10000 cells per well. Following the corresponding treatments, cell media was replaced with 30 μL of media containing 0.75 mM EU (Jena Bioscience, CLK-N002) and cells were incubated at 37°C for 30 min. After EU labeling, cells were fixed with 3.7% formaldehyde in PBS for 45 min at room temperature and permeabilized with 0.5% Triton X-100 in PBS for 30 min. Click-It reaction was performed using click-it buffer mix (100 mM Tris-HCl pH 8.5, 4 mM CuSO₄, 5 μM Alexa Fluor 488 azide, and 100 mM ascorbic acid added last and freshly prepared from solid powder) and incubating plate for 2 hr at room temperature in the dark. Wells were washed three times with 100 mM Tris-HCl pH 7.5, and nuclei were stained with 2 μg/mL DAPI in PBS for 30 min at room temperature in the dark. Cells were washed once with PBS, kept in PBS and visualized using High-throughput content screening (HCS) inverted widefield microscope from Olympus, with ScanR Acquisition and Analysis V2.8 software.

### DSK2 ubiquitin pull down

Isolation of poly-ubiquitylated RPB1 was performed as previously described ^76^. GST-DSK2 bead preparation and DSK2 pull down were conducted exactly as reported in ^77^. Briefly, cells were seeded with (tetracycline-free) FBS containing media in one 15 cm dish per condition. Cells treated with 20J/m^2^ and incubated at 37°C for 45 min after UV treatment were used as positive control. TENT lysis buffer (50 mM Tris-HCl pH7.4, 2 mM EDTA, 150 mM NaCl, 1 % (v/v) Triton X100), supplemented with freshly added 2 mM NEM and protease inhibitors were used for cell lysis. Samples were incubated in ice for 10 min, and 3 mM MgCl_2_ with 0.1% (v/v) DNARASE® were added, followed by rotation at 4°C for 1 hr. Extracts were collected from the supernatant, after 10 min centrifugation at 17000 g at 4° C, protein concentration was measured (Protein Assay Dye Reagent, Bio-Rad, 5000006) and adjusted to the same concentration (approximately 1mg) for each sample, at a final volume of 1 mL. Aliquots for input are taken at this step and mixed with SDS sample buffer; the remaining protein extracts are incubated with around 30 μL of packed GST-DSK2 beads overnight at 4° while rotating. Beads were collected with centrifugation (2500 rpm, 4° C) and washed twice with TENT buffer and once with PBS. Proteins are eluted by boiling beads in 80 µL 2xSDS sample buffer, followed by SDS-PAGE analysis.

### Quantitative PCR (qPCR)

For total RNA extraction of human samples growing in (tetracycline-free) FBS containing media, we used the RNeasy kit (QIAGEN, 74104) for mature RNA, following the instructions of the manufacturer including an on-column DNase treatment.

For HEK293 samples, reverse transcription was performed in 500 ng total RNA, using TaqMan Reverse Transcription Reagents (Thermo Fisher Scientific, N8080234). cDNA was amplified using HOT FIREPol® EvaGreen® qPCR Mix Plus (Solis BioDyne, 08-25-00001) with 40 cycles of 15 sec denaturation at 95°C, 20 sec annealing at 60°C, and 20 sec extension at 72 °C (melt curve included in the end).

For mouse samples, reverse transcription was performed in 500 ng total RNA, using SuperScript™ III Reverse Transcriptase (Invitrogen™, 18080093) together with RNaseOUT™ Recombinant Ribonuclease Inhibitor (Invitrogen™, 10777019). cDNA was amplified using PowerUp™ SYBR™ Green Master Mix for qPCR (Thermo Fisher Scientific, A25918) with a pre-step of 2 min at 50°C and initiation denaturation of 2 min at 95°C, followed by 40 cycles of 15 sec denaturation at 95°C, 1 min annealing/extension at 60°C (melt curve included in the end).

### polyA+ mRNA-seq in HEK293 cells

2×10^5^ cells were seeded in triplicates in (tetracycline-free) FBS containing media, in the presence or absence of 0.1 µg/µL doxycycline, in 6-well plates and were incubated at 37°C for 48 hrs.

For polyA+ mRNA-seq performed at SCAF1 KO and rescue cell lines, total RNA was extracted using Quick-RNA Microprep Kit (Zymo Research, R1051), following the instructions of the manufacturer including an on-column DNase treatment. Extracted RNA was measured with nanodrop and quality checked with Tapestation and 500 ng of RNA were sent to BGI genomics (https://www.bgi.com/global) for library preparation and sequencing. Samples were quenched with ∼50 million reads per sample.

For polyA+ mRNA-seq presented in Fig. 4 and Supplementary Fig. 4, cells were lysed in Trizol and total RNA was isolated with chloroform extraction followed by isopropanol precipitation. Libraries we prepared with 300 ng of RNA based on NEBNext Ultra II Directional RNA library prep kit for Illumina (NEB E7765) with NEBNext multiplex adapters for dual index UMI adapters (NEB E7416). Libraries were paired end sequenced with 55 reads in NextSeq 2000 (P3 100), with ∼67 million reads per sample.

### TT_chem_-seq

We performed TT_chem_-seq and as previously described ^39^. Briefly, two individual clones of *SCAF1* KO and rescue were seeded in (tetracycline-free) FBS containing media in duplicates and grown with and without doxycycline (final concentration 0.1 μg/µL) to cell confluency 80%, followed by exactly 15 min in vivo labelling with 1 mM 4SU (Glentham Life Sciences, GN6085). Cells were harvested with TRIzol and RNA was extracted accordingly to the manufacturer’s instructions. 4-thiouracil (4TU)-labeled RNA from S. *cerevisiae* (strain W303a) was added to each sample as spike in. The rest of the procedure was performed exactly as described in ^39^. 4SU-labbled RNA samples were measured with Qubit and fragmentation was confirmed by TapeStation (RNA HS assay). Libraries were prepared with 50 ng RNA, based on NEBNext Ultra II Directional RNA library prep kit for Illumina (NEB E7765-protocol for rRNA depleted FFPE RNA) with NEBNext multiplex adapters for dual index UMI adapters (NEB E7416). Libraries were single end sequenced with 61 reads in NextSeq 2000 (P3 50), with ∼47 million reads per sample.

### 4sUmRNA-seq in HEK293 and mouse cells

For HEK293 cells, two individual clones of *SCAF1* KO and rescue were seeded in (tetracycline-free) FBS containing media in duplicates and grown with and without doxycycline (final concentration 0.1 μg/µL) to cell confluency 80%, after 48 hrs. For mouse cells, one clone of the SCAF1 degron cell line was used and 2 hrs prior to 4sU addition, Auxin (IAA) was supplemented to half of the cells, seeded in triplicates.

Each sample was labelled for 1 hr with 1 mM 4SU (Glentham Life Sciences, GN6085), followed by TRIzol/chloroform extraction and isopropanol precipitation. 100 µg of RNA were mixed with 1 µg of 4-thiouracil (4TU)-labeled RNA from S. *cerevisiae* (strain W303a) and 3 µL of BB buffer (2.5 μL of 1 M Tris-HCl pH 7.4 and 0.5 μL of 0.5 M EDTA) in a total volume of 200 µL. Samples were biotinylated with the addition of 50 μL of 1 mg/mL EZ-Link HPDP-Biotin (Thermo Fisher, Cat. No. 21341; dissolved in DMF) and were incubated at room temperature for 3 hrs in the dark. RNA was purified with phenol:chloroform:isoamyl alcohol (25:24:1) and isopropanol precipitation. 4sU-biotinylated RNA was isolated using μMACS Streptavidin MicroBeads and columns equilibrated with nucleic acid equilibration buffer (Miltenyi Biotec, 130-074-101), following the manufacturer’s instructions. Libraries were prepared with 300 ng RNA, based on NEBNext Ultra II Directional RNA library prep kit for Illumina (NEB E7765 for polyA+ selection) with NEBNext multiplex adapters for dual index UMI adapters (NEB E7416). Libraries were paired end sequenced with 151 reads in P2 300 with ∼67 million reads per sample for HEK293 cells, and 55 reads P3 100 with ∼45 million reads per sample for mouse samples.

### Protein expression and purification

SCAF1-SRI domain with a C-terminal 6xHis-tag was gene synthesized (GenScript) and subcloned into a pHisSumo backbone (gift from Thomas Miller group) in the BamHI/XhoI sites. Wildtype SRI plasmids were transformed into Rosetta™ 2 *E. coli* competent cells. Liquid cultures were grown in 1 L LB media at 37°C to OD600 of 0.6-0.8 and induced with 0.4 mM IPTG for 4 hrs. Cells were harvested by centrifugation for 30 min (4000 rpm, 4° C) and resuspended in 15 mL lysis buffer (25 mM HEPES KOH pH=7.8, 300 mM NaCl, 10 mM imidazole, 1X Roche EDTA-free Complete protease inhibitor cocktail). The lysate was sonicated, followed by high-speed centrifugation for 20 min (30000 g, 4° C), and the supernatant was incubated for 15 min with 2 mL of Ni-NTA Superflow affinity resin (Qiagen) pre-equilibrated with lysis buffer. The resin was washed twice using gravity flow with 20 mL wash buffer (25 mM HEPES KOH pH 7.8, 300 mM NaCl, 20 mM imidazole, 1X Roche EDTA-free Complete protease inhibitor cocktail) and subsequently eluted with 250 mM imidazole in four fractions of 5 mL. Fractions containing the expressed protein were pooled, diluted 2.5x fold and the SUMO tag digested with 150 μg SUMO protease 1 (ULP1). The digestion was performed during dialysis against buffer A (25 mM HEPES KOH pH 7.8, 150 mM NaCl, 1 mM DTT) at 4°C overnight. Digested proteins were centrifuged for 10 min (4000xg, 4° C) and loaded onto an equilibrated HiTrap (Cytiva) 1 mL SP HP column using the ÄKTA Pure FPLC system. The column was washed, and SCAF1-SRI was separated from the protease and cleaved SUMO-tag with a gradient to 100% buffer B (25 mM HEPES KOH pH 7.8, 1 M NaCl, 1 mM DTT). Fractions containing SCAF1-SRI were concentrated to 300 μL using the 10,000 MWCO centrifugal filters (Amicon). The buffer was exchanged to 150 mM final NaCl concentration, and the sample was aliquoted, flash-frozen (liquid nitrogen) and stored at - 20°C.

### Fluorescence polarization (FP) assay

Carboxyfluorescein (FAM)-labeled peptides representing the unphosphorylated, single phosphorylated and double phosphorylated RNAII CTD phospho-peptides functioned as probes for quantifying the interaction between the RNAPII CTD and SCAF1-SRI. The probes were diluted in Exchange buffer (50 mM HEPES, pH 7.4, 400 mM NaCl, 0.025% DDM, 10% (v/v) glycerol) and 10 µL of each was added to a final concentration of 250 nM per well in a black 384-well plate (Corning). The SCAF1-SRI protein (starting concentration of 20 µM) was diluted using a two-fold dilution series; 30 µL of the SCAF1-SRI was added to each well to final concentrations ranging from 0.305 nM to 5000 nM. Blanks containing 10 µL probe and 30 µL Exchange buffer were included. The plate was spun down at 1000 rpm for 1 min and the FP signal was measured on a Flex station using SoftMax Pro software 7.0. Each experiment was conducted in duplicate. The binding curves were generated by plotting the FP signal, expressed as millipolarization units (mP), against SCAF1-SRI protein concentration for each probe included.

### polyA+ mRNA-seq in mouse cells

Cells were differentiated prior to mRNA extraction, as described in the corresponding sections. Total RNA was isolated using the RNeasy kit (QIAGEN, 74104), following the instructions of the manufacturer including an on-column DNase treatment. Libraries were prepared with 300 ng of RNA based on NEBNext Ultra II Directional RNA library prep kit for Illumina (NEB E7765, for polyA+ selection) with NEBNext multiplex adapters for dual index UMI adapters (NEB E7416). Libraries were paired end sequenced with 55 reads in NextSeq 2000 (in P3 100 or P2 100), with ∼33 million and ∼42 million reads per sample (for non-direct lineage differentiation and neuronal differentiation respectively).

## QUANTIFICATION AND STATISTICAL ANALYSIS

### Sequencing data processing and bigwig file generation

The quality of raw sequencing reads from polyA+ mRNA-seq in HEK293 cells was reads checked using FastQC (v0.11.9) (Babraham Bioinformatics). Sequencing data of TT_chem_-seq, 4sU mRNA-seq and polyA+mRNA-seq in mouse cells were demultiplexed using Illumina bcl2fastq and then reads quality was checked with fastQC We sequenced the UMI sequences and the DNA fragments in two separate cycles, generating one (single-end reads) or two (paired-end reads) DNA sequence read/reads with an extra UMI sequence read. The UMI sequences were then extracted and attached to the headers of the DNA read sequences using UMI-tools ^78^.

The universal Illumina sequencing adapters were trimmed using trim_galore (https://www.bioinformatics.babraham.ac.uk/projects/trim_galore/) and cutadapt ^79^ with default setting.

Reads were mapped to a merged reference of the human or mouse genome (Gencode: GRCh38.primary_assembly.genome.fa for human and GRCm39.genome.fa, M35 release for mouse) and yeast genome (Ensembl: Saccharomyces_cerevisiae.R64-1-1.dna.toplevel.fa) using STAR (2.7.9a) ^80^ with options: -- runThreadN 10 --outSAMattributes NH HI AS NM MD XS --outSAMstrandField intronMotif -- outSAMmultNmax 1 --outSAMunmapped None --outSAMtype BAM. Uniquely mapped reads were extracted from the aligned bam files with samtools (v1.15.1) ^81^. PCR duplicates were identified and removed with UMI-tools ^78^. The uniquely mapped reads were converted to bedgraphs (with spike in or library size normalized) and then to bigwig format using bedtools (2.30.0) ^82^ and GenomeToolset.

### Downstream analysis of mRNA-seq and 4sUmRNA-seq

Genes that were uniquely assigned to each feature were counted using featureCounts function from Subread. The raw count tables of gene features were used to identify differentially expressed genes between *SCAF1* KO (or degradation) and rescue (or WT) using DESeq2 ^83^. The reads that mapped to each exon were counted using DEXseq from HTSeq (v2.0.2) ^37^, with the count table of exon features were used to identify differentially used exons between *SCAF1* KO (or degradation) and rescue (or WT). The alternative splicing event changes between *SCAF1* KO and rescue were analyzed using rMATS ^36^, with default settings.

### APAlyzer analysis

To assess changes in alternative polyadenylation (APA) between conditions, we used APAlyzer package (v1.22.0) ^38^, with mRNA-seq data mapped to custom made reference intronic polyadenylation sites, using the reference gtf files (GRCh38 for human and GRCm39 for mouse), as described at ^38^. Default setting were used for the remaining analysis.

### Metagene profiling from TT_chem_-seq data

To explore the coverage of metagenes (i.e. mean coverage across genes), we used Easeq software ^84^ for non-overlapping protein-coding and lncRNA genes and for genes that showed increased intronic polyA usage (APAlyzer analysis) for the regions around transcription start site (TSS), gene bodies and transcription end site (TES). For gene body, we plotted the region from 10kb upstream of TSS to 10kb downstream of TES. The coverage between TSS and TES for gene body were scaled into 100 bins for each gene. For TSS and TES, we calculated the coverage 4kb upstream and 4kb downstream for each single nucleotide position. Then we averaged the coverages for genes transcribed from both the sense and antisense strands. Visualization and preceding analysis was done using EaSeq and its integrated tool Easeq software ^84^.

### Analysis of CUT & RUN data

FASTQ data from Cut & Run sequencing performed in ^41^ were checked with FASTQC and were analyzed as described in the ^41^. Briefly, reads were mapped using Bowtie2 (2.5.3) ^85^ to merged genome of ensemble acquired human (GRCh38) and Escherichia coli (Escherichia_coli_str_k_12_substr_mg1655_gca_000005845.ASM584v2.dna.toplevel.fa).

Uniquely mapped reads were extracted from the aligned bam files with samtools (v1.15.1) ^81^, were spike in normalized and converted to bedgraph files with bedtools (2.30.0). Replicates were averaged and metagene analysis was performed as described at metagene profiling from TT_chem_-seq data. Visualization of the data was performed using Easeq ^84^.

## Supplemental item titles

Table S1: Mass spectrometry of SCAF1 and RNAPII immunoprecipitation experiments

Table S2: List of oligonucleotides

## Acknowledgments

This work was supported by grants to L.H.G from the European Research Council (ERC Agreement, TranscriptStress, 101076758), a Hallas-Møller emerging investigator grant to from the Novo Nordisk Foundation (NNF20OC0059959), and a Sapere Aude research leader program grant from Independent Research Fund Denmark (0165-00092B). Research in the Center for Gene Expression (CGEN) is funded by the Danish National Research Foundation (DNRF166). G.M.H is funded by Læge Sofus Carl Emil Friis and Hustru Olga Doris Friis’ legat. E.K. is supported by a Lundbeckfonden (Lundbeck Foundation) grant (R380-2021-1519). Work in the group of T.C.R.M was supported by a Novo Nordisk Fonden Hallas-Møller Emerging Investigator Grant (NNF22OC0073571), the Danish National Research Foundation (DNRF115) and Carlsberg Foundation grant CF23-0803. Work in the group of J.J.Z. was supported by a Hallas-Møller emerging investigator grant to from the Novo Nordisk Foundation (NNF21CC0073729) and a Sapere Aude research leader program grant from Independent Research Fund Denmark (0169-00031B). Mass spectrometry-based proteomics analyses were performed by the Proteomics Research Infrastructure (PRI) at the University of Copenhagen (UCPH), supported by the Novo Nordisk Foundation (NNF) (grant agreement number NNF19SA0059305). We thank the staff of the CPR/reNEW Genomics Platform for support: H. Wollmann, M.Michaut, A. Kalvisa, C. Emmerson. The Novo Nordisk Foundation Center for Stem Cell Medicine (reNEW) is supported by the Novo Nordisk Foundation grant number NNF21CC0073729. The Novo Nordisk Foundation Center for Protein Research (CPR) is supported by the Novo Nordisk Foundation grant number NNF14CC0001. Lastly we would like to thank Jesper Q. Svejstrup for feedback on the manuscript.

## Author contributions

L.H.G. and S.K. conceived the project. S.K. performed the main experiments, S.F.R assisted with SCAF1-SRI purification and G.M.H. performed SCAF1-SRI *in vitro* peptide binding assay. N.S generated the SCAF1 degron cell lines in mESCs and assisted mESCs culturing. N.S. and E.K. assisted with the differentiation protocol and validation. S.K. and H.L. performed the bioinformatic analysis. L.H.G, T.CR. M. and J.J.Z. supervised the project. S.K. and L.H.G. wrote the manuscript. All authors read and approved the final version of the manuscript.

