## Supplemental Figures for "SCAF1 driven polyadenylation site usage regulates mRNA isoform expression and neuronal differentiation"

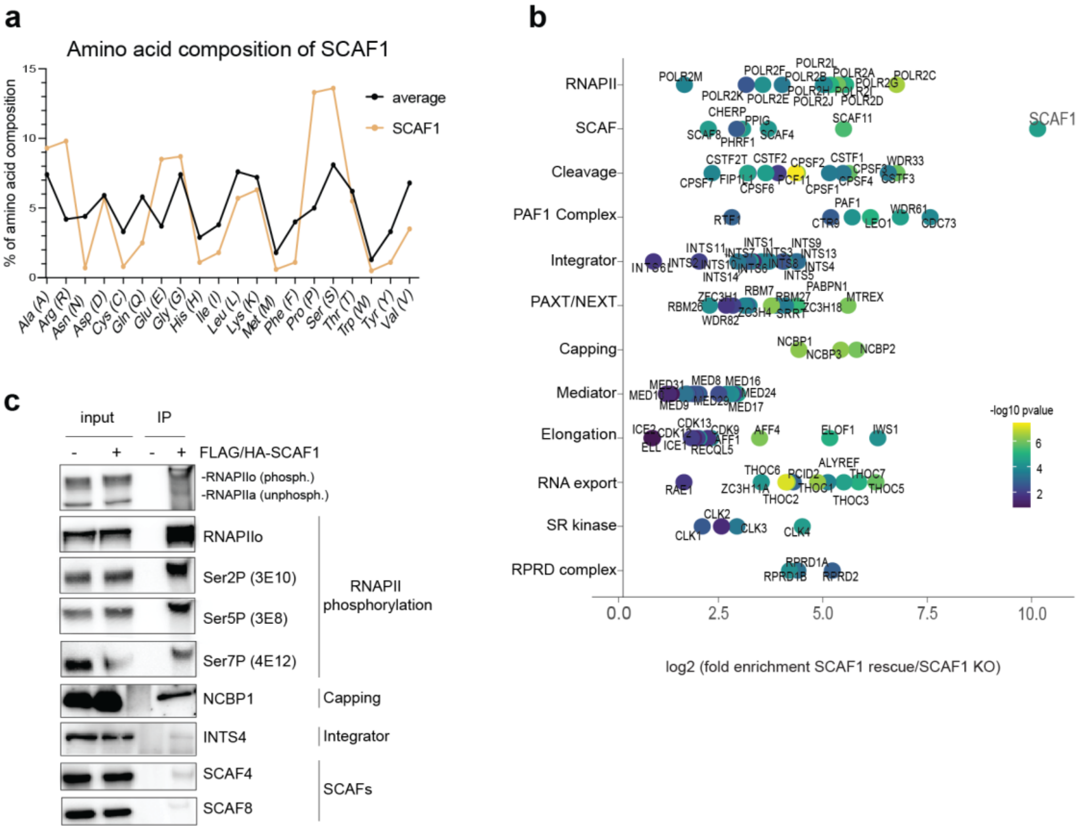
a
Amino acid composition of SCAF1
% of amino acid composition
average
SCAF1
Ala (A)
Arg (R)
Asn (N)
Asp (D)
Cys (C)
Gln (Q)
Glu (E)
Gly (G)
His (H)
Ile (I)
Leu (L)
Lys (K)
Met (M)
Phe (F)
Pro (P)
Ser (S)
Thr (T)
Trp (W)
Tyr (Y)
Val (V)
b
RNAPII
SCAF
Cleavage
PAF1 Complex
Integrator
PAXT/NEXT
Capping
Mediator
Elongation
RNA export
SR kinase
RPRD complex
SCAF1
-log10 pvalue
log2 (fold enrichment SCAF1 rescue/SCAF1 KO)
c
input
IP
FLAG/HA-SCAF1
-RNAPIIo (phosph.)
-RNAPIIa (unphosph.)
RNAPIIo
Ser2P (3E10)
Ser5P (3E8)
Ser7P (4E12)
RNAPII phosphorylation
NCBP1
Capping
INTS4
Integrator
SCAF4
SCAF8
SCAFs

Sup. Figure 1

***Supplementary Fig. 1: Further investigation on SCAF1 interactome.***

**a** Amino acid frequency of SCAF1, compared to average protein composition (based on http://www.nimbios.org/~gross/bioed/webmodules/aminoacid.htm). **b** Log2 fold enrichment of SCAF1 interacting proteins, grouped in different categories, based on the complexes they belong. Proteins are scaled colored according to the interaction significance (-log10 p-value > 1.3). **c** Western blot validation of enriched proteins in the FLAGHA-SCAF1 immunoprecipitates, compared to control, untagged wildtype cell lines.

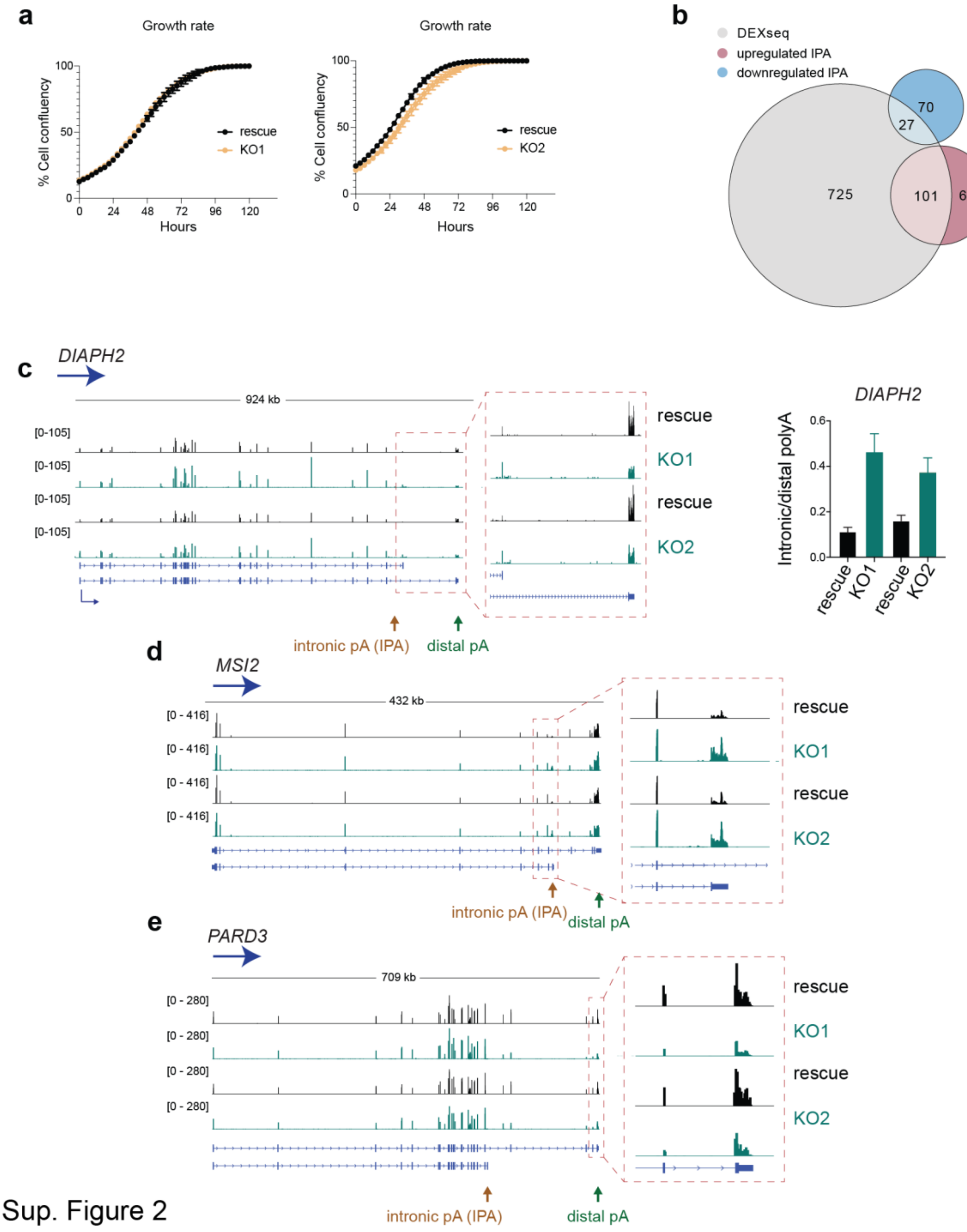


1287 Sup. Figure 2

***Supplementary Fig. 2: SCAF1 KO promotes alternative polyadenylation***

**a** Cell growth in rescue compared to *SCAF1* KO, in two clones, monitored by Incucyte. Error bars represent ±SD. **b** Venn diagram of the number of genes with differential exon usage (DEX-seq hits), of hits that show upregulated and downregulated usage of intronic APA sites, and their intersection. Gene overlap between DEXseq and IPA is statistically significant, as assessed using hypergeometric test (p-value <0.0001, ****). **c-e** IGV tracks of more gene examples, identified through APAlyzer analysis for upregulated usage of intronic APA sites of the mRNA-seq samples. Intronic and distal polyA sites pointed with orange and green arrows respectively. (**c**) Additional validation of proximal to distal mRNA isoform expression by qPCR with primer-pairs for *DIAPH2*. Primer-pairs used were designed to be specific for the short or the long isoform 3' UTR. The ratio of intronic to distal polyA was calculated as the ratio between products from the short to long isoform 3' UTR and the signal ratio was normalized to *GAPDH* and a gene specific reference region (common in both isoforms). Error bars represent ±SD.

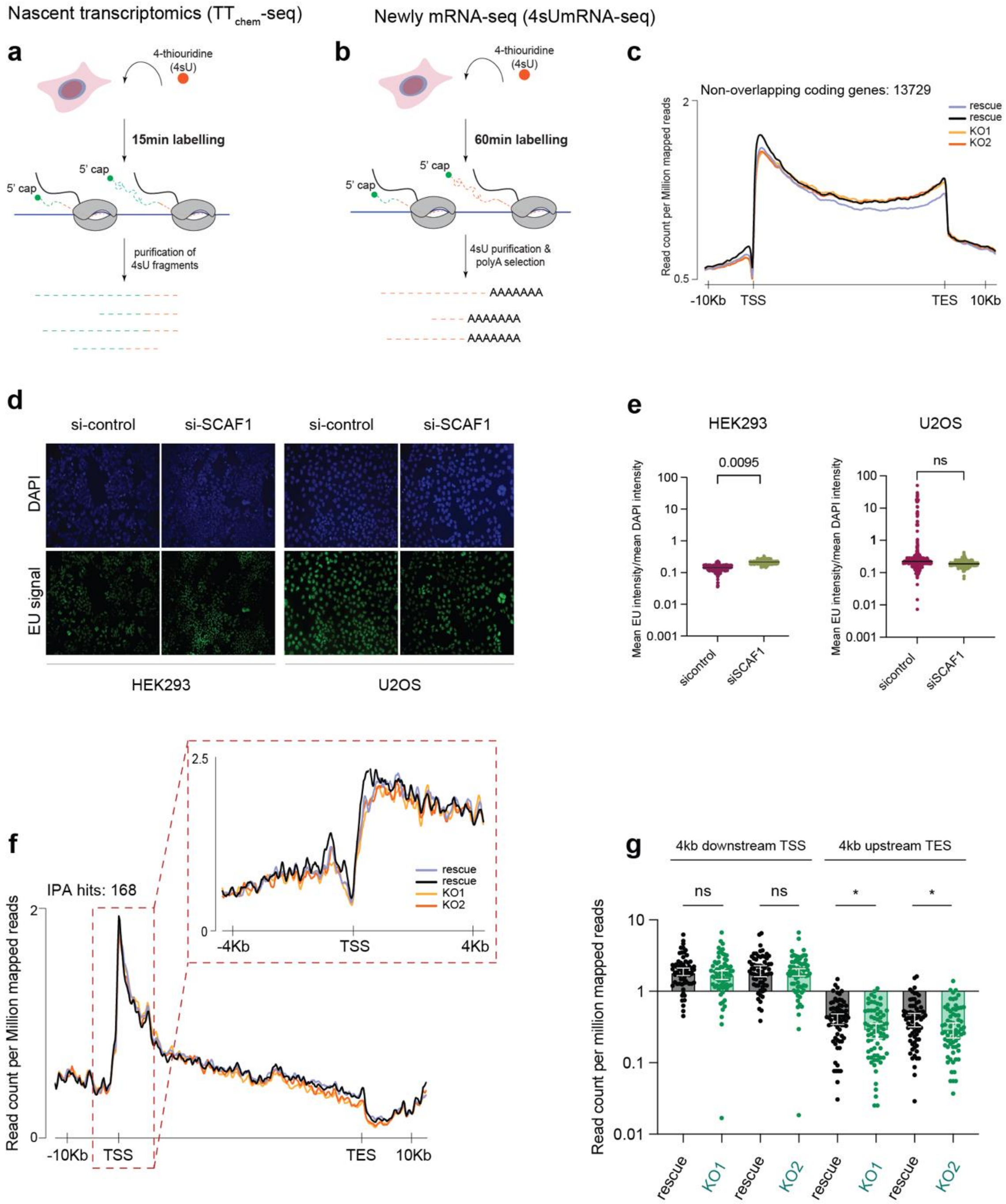
Nascent transcriptomics (TTchem-seq)
Newly mRNA-seq (4sUmRNA-seq)
a
4-thiouridine (4sU)
15min labelling
5' cap
5' cap
purification of 4sU fragments
b
4-thiouridine (4sU)
60min labelling
5' cap
5' cap
4sU purification & polyA selection
AAAAAAA
AAAAAAA
AAAAAAA
c
Non-overlapping coding genes: 13729
rescue
rescue
KO1
KO2
Read count per Million mapped reads
-10Kb
TSS
TES
10Kb
d
si-control
si-SCAF1
si-control
si-SCAF1
DAPI
EU signal
HEK293
U2OS
e
HEK293
U2OS
0.0095
ns
Mean EU intensity/mean DAPI intensity
sicontrol
siSCAF1
f
IPA hits: 168
Read count per Million mapped reads
-10Kb
TSS
TES
10Kb
-4Kb
TSS
4Kb
rescue
rescue
KO1
KO2
g
4kb downstream TSS
4kb upstream TES
ns
ns
*
*
Read count per million mapped reads
rescue
KO1
rescue
KO2
rescue
KO1
rescue
KO2

Sup. Figure 3

***Supplementary Fig. 3: SCAF1 regulates target genes co-transcriptionally without affecting transcription globally***

**a, b** Graphical outline of isolation of nascent transcriptome, $TT_{chem}$-seq (**a**), and of the newly synthesized full length (polyA selected) transcriptome with 4sUmRNA-seq (**b**). The different 4sU labelling times are highlighted in the two protocols. **c** Metagene plot of $TT_{chem}$-seq signal against all non-overlapping genes (13729), in the two *SCAF1* KO and their corresponding rescue clones. Signal resulted from duplicate merging of the samples, gene sizes are scale normalized with 10 kb region upstream and downstream of the genes plotted. Signal resulted from duplicate merging of the samples. **d** Images representing global nascent transcriptomic through EU labelling in HEK293 and U2OS, upon SCAF1 depletion (siRNA-SCAF1) and wildtype (siRNA-control) conditions. **e** Violin plots presenting EU signal quantification, normalized to background DAPI signal in wildtype and upon SCAF1 depletion, in the different cell lines shown in (**d**). Nested t-test performed for statistical significance, with p-value indicated in the plot. **f** Metagene profile of $TT_{chem}$-seq against the 168 target genes of SCAF1 that display increased usage of intronic APA sites, with zoom in 4 kb upstream and downstream of the transcription. **g** Read coverage quantification of the $TT_{chem}$-seq data presented in (**f**). Read count of the signal measured within a 4 kb window downstream from TTS, as well as read count signal within a 4 kb window upstream of TES, in RPKM, for the two *SCAF1* KO clones and their corresponding rescues. Calculations were performed using build in Easeq tools. Statistical significance measured by running mann-whitney unpaired t-test with p-value<0.0001, ****, with p-values indicated at the plot.

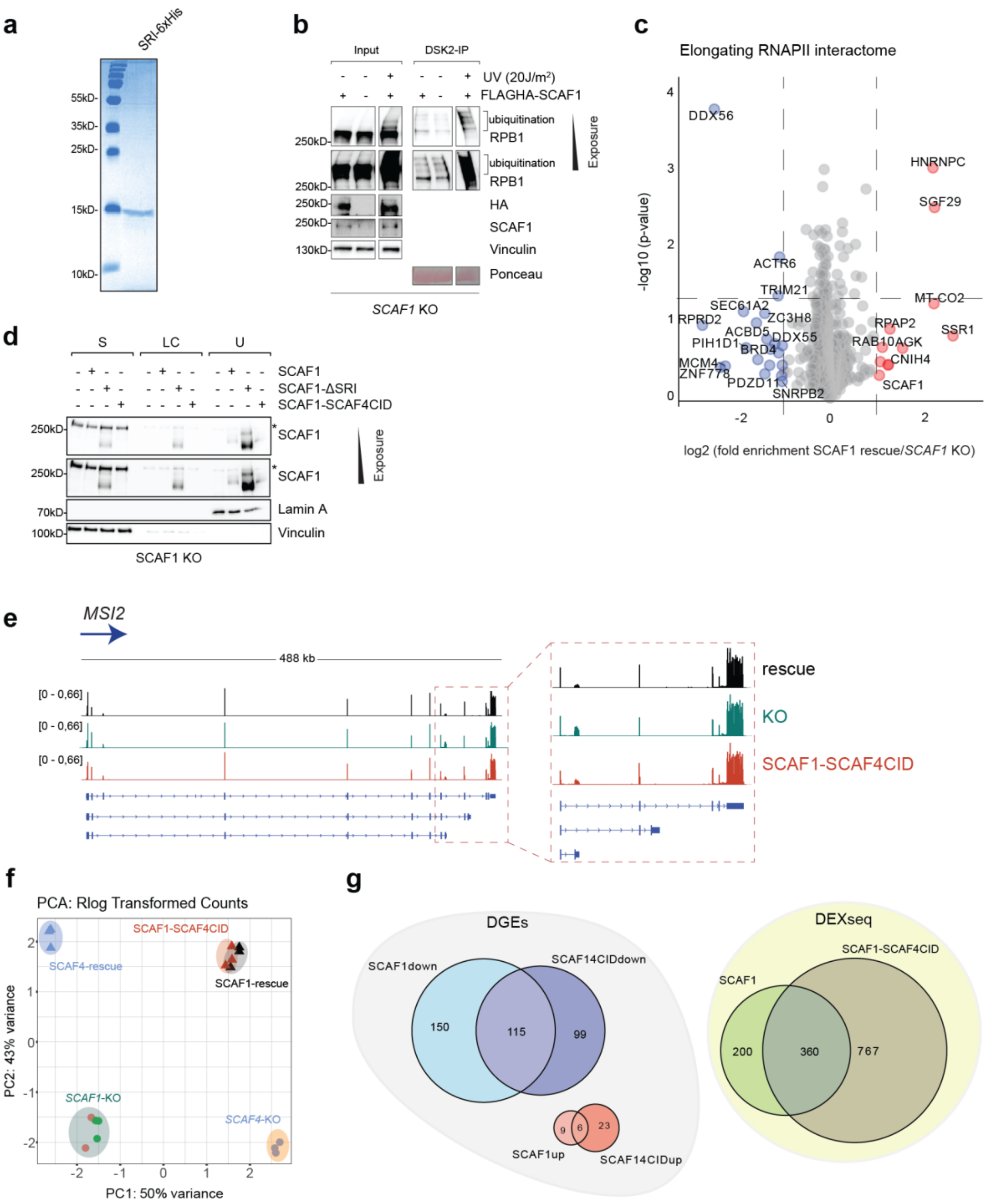


1326 Sup. Figure 4

***Supplementary Fig. 4: SCAF1 binding to Ser2/Ser5 double phosphorylated RNAPII is required for SCAF1 dependent phenotype***

**a** Coomassie stain verifying 6xHis-SRI purification. **b** Western blot of DSK2 pull-down in rescue and *SCAF1* KO cell lines. Rescue cells treated with 20J/m2 of UV were used as positive control. **c** Volcano plot showing the differentially associating proteins to RNAPII, in rescue and *SCAF1* KO background. Proteins with reduced RNAPII association upon *SCAF1* KO are blue colored, whereas red dots mark the proteins that increase interaction with RNAPII, upon *SCAF1* KO. 4H8 antibody, recognizing the N-terminus of Rpb1, was used for precipitating elongating RNAPII. Vertical dashed lines mark log2FC=-1 and log2FC=1, and the horizontal one indicates significantly altered interacting proteins, with -log10p-value =1.3 (for p-value=0.05). **d** Western blot of cellular fractions (cytoplasmic: **S**, low chromatin binding: **LC**, High stringency buffer Urea: **U**) from SCAF1 rescue, SCAF1-ΔSRI and SCAF1-SCAF4CID samples. LaminA and vinculin are used as loading and fractionation controls. Two different exposures of SCAF1 blots are shown and asterisk indicates the presence of non-specific band. **e** IGV tracks, in support to Fig. 4e-f examples for differential expression of the full length (green arrow) and short (orange arrow) isoforms from the mRNA-seq data of SCAF1 rescue, *SCAF1* KO and *SCAF1-SCAF4CID* cell lines. **f** Principal component analysis (PCA) plot of the different cell lines that cluster, based on mRNA-seq data. Cell lines are distinguished by different color, with each point represents an individual replicate (total number of replicates=3 per cell lines). SCAF4 rescue and *SCAF4* KO, generated at [21] are used as control of SCAF1-SCAF4CID. **g** Venn diagram illustrating the overlap between targets of SCAF1 rescue and SCAF1-SCAF4CID rescue. Comparisons of the differentially expressed genes (DGEs) and genes that show differential exon usage (DEX-seq) between **SCAF1 rescue** and **SCAF1-SCAF4CID rescue** cell lines, both compared to *SCAF1* KO, and their overlapping targets.

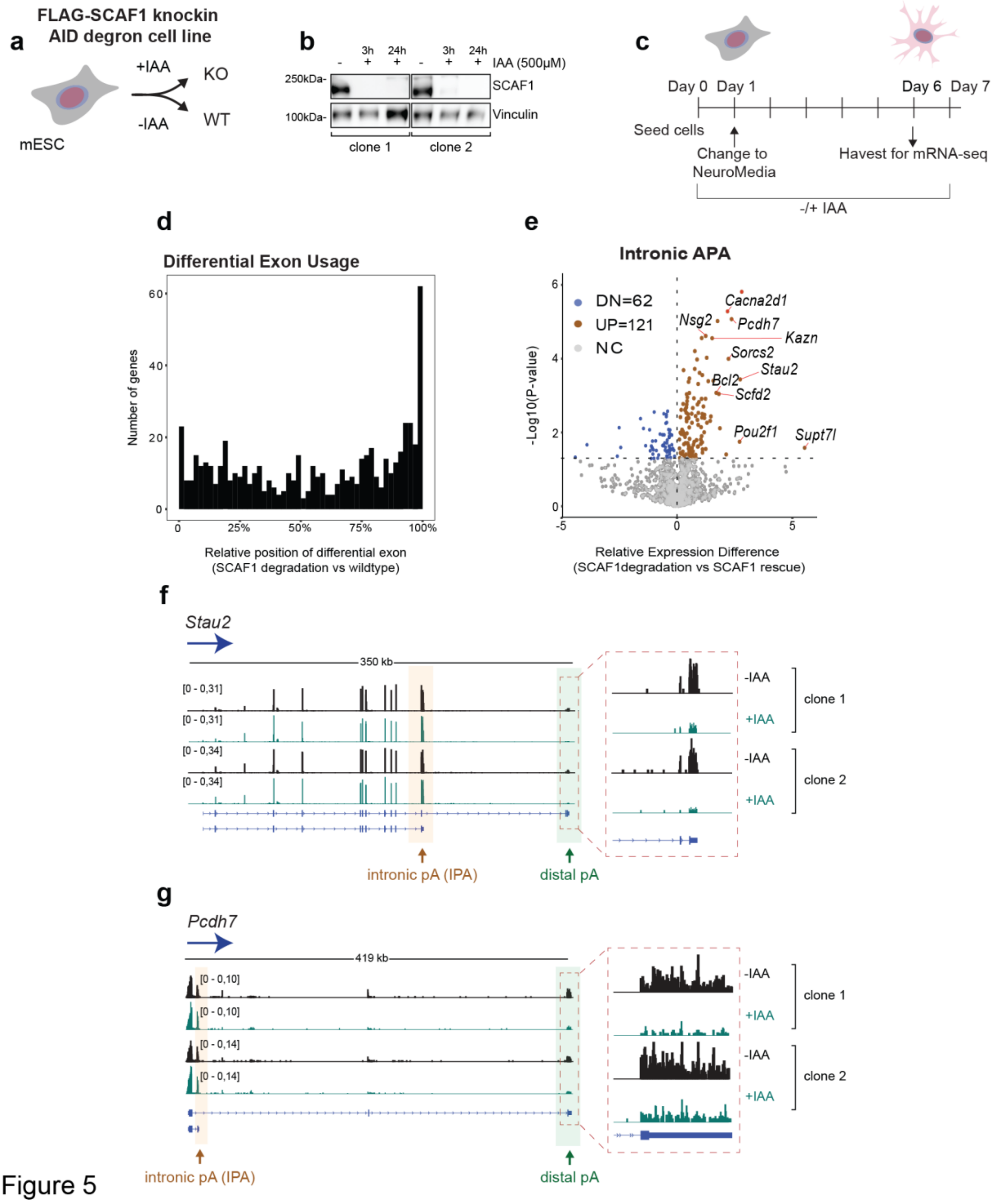
a
FLAG-SCAF1 knockin
AID degron cell line
+IAA
KO
-IAA
WT
mESC
b
3h
24h
3h
24h
IAA (500μM)
250kDa-
SCAF1
100kDa-
Vinculin
clone 1
clone 2
c
Day 0 Day 1
Day 6 Day 7
Seed cells
Change to NeuroMedia
Havest for mRNA-seq
-/+ IAA
d
Differential Exon Usage
Number of genes
Relative position of differential exon
(SCAF1 degradation vs wildtype)
e
Intronic APA
DN=62
UP=121
NC
Cacna2d1
Nsg2
Pcdh7
Kazn
Sorcs2
Stau2
Bcl2
Scfd2
Pou2f1
Supt7l
-Log10(P-value)
Relative Expression Difference
(SCAF1degradation vs SCAF1 rescue)
f
Stau2
350 kb
[0 - 0,31]
[0 - 0,34]
-IAA
+IAA
clone 1
clone 2
intronic pA (IPA)
distal pA
g
Pcdh7
419 kb
[0 - 0,10]
[0 - 0,14]
-IAA
+IAA
clone 1
clone 2
intronic pA (IPA)
distal pA

Figure 5

***Supplementary Fig. 5: SCAF1 extensive degradation shows conserved transcriptional phenotype in mouse cells and plays a potential role during neuronal differentiation***

**a, b, c** Analysis of 4sUmRNA-seq data from 3 hrs SCAF1degradation in mESCs in naïve conditions. (**a**) Schematic outline of the samples collected for 4sUmRNA-seq, in wildtype and 3h SCAF1 degradation. (**b**) Principal component analysis (PCA) plot of the SCAF1 degron mESCs (+/-IAA) 4sUmRNA-seq. One SCAF1 degron cell lines is used and each point represents an individual replicate (total number of replicates=3). (**c**) Heatmap from 1h-mRNA-seq data of SCAF1 degron cells. The four differentially expressed genes from 4sUmRNA-seq analysis are displayed, with log2 fold change illustrated in scaled color. **d** Barplot of the genes with differential exon usage, identified through DEX-seq analysis of 3h SCAF1 degradation compared to wildtype. **e** Schematic representation depicting the experimental workflow. 1) **Naïve** SCAF1 degron mESCs, growing in 2iLiF conditions were subjected to: 2 days in 2iLiF (+/-IAA), differentiation to 2) **EpiLCs** for 2 days (+/-IAA) according to [74] protocol, or exit from pluripotency and default 3) **Differentiation** for 2 days upon 2iLiF withdrawal (+/-IAA). **f** Hierarchical clustering and heatmap of sample-to-sample distances using variance-stabilized (vst) transformed read counts. Clustering was performed across samples and was visualized as a dendrogram. Color intensity representing the degree of dissimilarity between samples. **g** Bubble plot illustrating the fold enrichment of Gene Ontology (GO) biological processes associated with genes differentially expressed in SCAF1 degron mESCs upon 2iLiF withdrawal with or without IAA. Color intensity indicates the statistical significance (–log10 p-value), while bubble size represents the number of genes linked to each GO term. **h** Venn diagram of the number of DEX-seq hits, of hits that show increased usage of intronic APA, and their intersection from mRNA-seq analysis of the SCAF1 degron neuronal differentiated mESCs (day 6) **i** Venn diagram of the number of the hits showing increased usage of intronic APA in human and mouse, and their intersection. Overlapped genes, also colored green, are statistically significant, as assessed with hypergeometric test (p-value cutoff = 0.01. Overall overlap significance for the 25 genes, with p-value= 2.397188e-18: p-value= 2.357391e-16 for the upregulated IPA and p-value= 0.0002337947 for the downregulated).

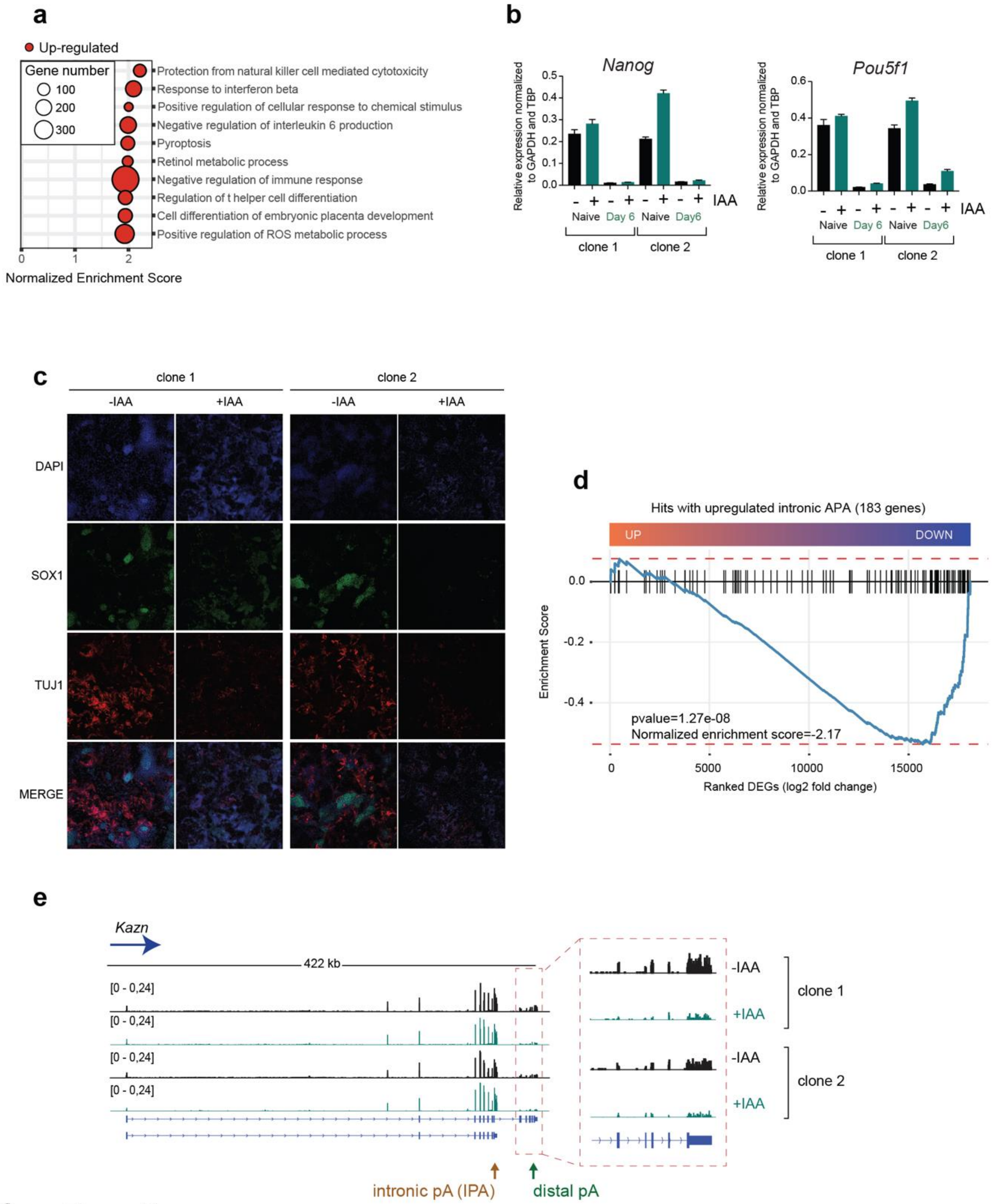
a
Up-regulated
Gene number
100
200
300
Protection from natural killer cell mediated cytotoxicity
Response to interferon beta
Positive regulation of cellular response to chemical stimulus
Negative regulation of interleukin 6 production
Pyroptosis
Retinol metabolic process
Negative regulation of immune response
Regulation of t helper cell differentiation
Cell differentiation of embryonic placenta development
Positive regulation of ROS metabolic process
Normalized Enrichment Score
b
Nanog
Pou5f1
Relative expression normalized to GAPDH and TBP
IAA
Naive
Day 6
Day6
clone 1
clone 2
c
clone 1
clone 2
-IAA
+IAA
DAPI
SOX1
TUJ1
MERGE
d
Hits with upregulated intronic APA (183 genes)
UP
DOWN
Enrichment Score
pvalue=1.27e-08
Normalized enrichment score=-2.17
Ranked DEGs (log2 fold change)
e
Kazn
422 kb
[0 - 0,24]
-IAA
+IAA
clone 1
clone 2
intronic pA (IPA)
distal pA

Sup. Figure 6

***Supplementary Fig. 6: Loss of SCAF1 impairs neuronal differentiation***

**a** Bubble plot from GSEA of the expression changes (SCAF1 degradation vs wildtype). The top 10 most significantly upregulated pathways are shown, with normalized enrichment scores (padj < 0.05) and bubble sizes are scaled accordingly to number of upregulated genes within individual categories. **b** qPCR validation of the expression pattern of two pluripotency genes in the two SCAF1 degron clones tested in naïve and neuronal differentiated samples. *Gapdh* and *Tbp* were used for expression normalization and error bars represent ±SD. **c** IF images captured at 10x magnification, using the nuclear, neuronal stem cell marker SOX1 and the marker for early neurons, TUJ1, in the two SCAF1 degron clones. The experiment was performed twice. **d** Pre-ranked gene set enrichment analysis of mRNA-seq (x-axis), from the most upregulated to the most downregulated genes identified from DESeq2 analysis. Blue line indicates the trend of the genes with upregulated intronic APA in differentiated mouse cells in regard to differential expressed genes. Statistical significance (p-value) and normalized enrichment score are indicated. **e** Representative gene example from *Kazn* gene that regulates neuronal development and was identified also via APAlyzer analysis, showing differential expression of the full length and the short version of the corresponding transcripts in the two SCAF1 degron cell lines. Orange and green arrows point to the annotated polyA sites, proximal and distal respectively.
